# Intracranial Targeting of Cholesterol Processing Reveals a Therapeutic Vulnerability that Reprograms Glioblastoma and Promotes Antitumor Immunity

**DOI:** 10.64898/2026.08.18.745568

**Authors:** Maria J. Ulloa-Navas, Rachel M. Whitehead, Vanessa K. Jones, Michaelides Loizos, Mieu Brooks, Aleeshba N. Basil, Raquel Morales-Gallel, Cristina Gomez-Palmero, G. Abril Reynaga-Macias, Jesus E. Sanchez-Garavito, Brianna Tapia-Dierking, Asha A. Nair, Juan P. Navarro-Garcia de Llano, Paula Schiapparelli, Ian Dryden, Steven Rosenfeld, Victoria E. Clark, Haidong Dong, Loic Deleyrolle, Hong Qin, Vicente Herranz-Perez, Yingxue Ren, Jose M. Garcia-Verdugo, Alfredo Quinones-Hinojosa

**Affiliations:** Department of Neurosurgery, Mayo Clinic, Jacksonville, FL, USA; Icahn School of Medicine, New York, NY USA; Laboratory of Comparative Neurobiology, University of Valencia-CIBERNED, Valencia, Spain; Department of Quantitative Health Sciences, Mayo Clinic, Jacksonville, FL, USA; Department of Pathology, Mayo Clinic, Jacksonville, FL, USA; Department of Neurology, Mayo Clinic, Jacksonville, FL, USA; Department of Urology, Mayo Clinic, Rochester, MN, USA; Department of Cancer Biology, Mayo Clinic, Jacksonville, FL, USA; Department of Hematology-Oncology, Mayo Clinic, Jacksonville, FL, USA

**Author notes:** Corresponding Author: Alfredo Quinones-Hinojosa 4500, San Pablo Rd S, Mayo Clinic, Jacksonville, FL. Author Passed Away during the preparation on this manuscript (Jose M. Garcia-Verdugo).

**Keywords:** Glioblastoma, cholesterol metabolism, ER stress, unfolded protein response, autophagy, metabolic therapy, clemastine, bexarotene, intracranial delivery

## Abstract

Glioblastoma (GBM) remains the most lethal primary brain cancer due to its remarkable metabolic plasticity and therapeutic resistance. Here, we identify cholesterol dependency as a therapeutically exploitable vulnerability in GBM using two FDA-approved drugs: the H1 histamine antagonist clemastine and the retinoid X receptor agonist bexarotene. Combined treatment induces potent synergistic anti-tumor activity across patient-derived glioma models, suppressing proliferation, stemness, and survival at sub-IC50 concentrations. Mechanistically, this therapy disrupts cholesterol biosynthesis, transport, and homeostasis, triggering endoplasmic reticulum stress and activation of the unfolded protein response, ultimately leading to autophagy and apoptotic cell death. Orthotopic patient-derived glioma models recapitulate these mechanisms in vivo, where local intracranial administration significantly reduces tumor progression and prolongs survival using fourfold lower doses than systemic intraperitoneal delivery. Single-cell RNA sequencing revealed activation of regeneration and plasticity programs, accompanied by immune microenvironment remodeling and enhanced inflammatory signaling. Importantly, syngeneic models preserved immune cell composition, supporting future integration with immunotherapeutic strategies. Together, these findings establish cholesterol dysregulation–induced metabolic collapse as a promising therapeutic approach for GBM.

## INTRODUCTION

Glioblastoma (GBM), a primary brain cancer, remains among the most lethal human malignancies, with a median overall survival of approximately 15 months despite maximal safe resection followed by radiotherapy and temozolomide chemotherapy^1^. A defining feature of GBM is its capacity to adapt to therapeutic and environmental stress through metabolic reprogramming, thereby promoting tumor persistence, near-universal recurrence, and resistance to therapy^2–4^. While genomic heterogeneity has historically dominated the conceptual framework of GBM biology, emerging evidence suggests that metabolic plasticity represents a fundamental and targetable driver of tumor survival. Single-cell studies have further shown that malignant GBM cells dynamically occupy astrocyte-like, oligodendrocyte progenitor-like, neural progenitor-like, and mesenchymal-like states that are shaped by both genetic and microenvironmental cues^3^. In particular, glioma stem cells (GSCs) exhibit pronounced dependence on lipid and cholesterol metabolism to sustain cell division, membrane biogenesis, oncogenic signaling, oxidative stress buffering, and adaptation to the nutrient- and oxygen-limited brain tumor microenvironment ^2,4,5^. However, how cholesterol dependency can be therapeutically exploited to induce irreversible metabolic collapse in GBM without compromising normal neural and glial homeostasis remains incompletely understood.

Cholesterol homeostasis is tightly regulated through coordinated control of biosynthesis, uptake, trafficking, storage, esterification, and efflux ^5^. Unlike many peripheral cancers that rely heavily on de novo cholesterol biosynthesis, GBM cells preferentially obtain cholesterol from the brain microenvironment through active uptake and scavenging pathways ^4^. In the central nervous system, astrocytes are major suppliers of extracellular cholesterol, and brain tumors evolve within a lipid-rich milieu shaped by extensive myelin turnover and myeloid cell activity^4,6,7^. Consequently, GBM cells upregulate sterol uptake and transport programs together with lipid droplet formation to sustain rapid proliferation and survive metabolic stress, whereas de novo cholesterol biosynthesis appears to function more as a limited compensatory pathway^4,8^. Lipid droplets act as dynamic reservoirs that buffer excess fatty acids and cholesterol esters, thereby protecting cells from lipotoxicity and membrane instability^8^. Perturbation of these pathways can alter endoplasmic reticulum (ER) membrane composition and protein-folding capacity, leading to ER stress and activation of the unfolded protein response (UPR). The UPR is mediated by the ER stress sensors PERK, IRE1, and ATF6, which collectively attempt to restore proteostasis by reducing protein translation, increasing chaperone activity, and enhancing degradation of damaged proteins^9^. While transient UPR activation may initially support tumor adaptation and survival, sustained ER stress can instead drive chronic proteotoxic stress, autophagy, and apoptotic cell death^9^. Increasing evidence further links persistent lipid imbalance and aberrant lipid droplet accumulation to ER stress vulnerability, directly connecting cholesterol dysregulation to a targetable metabolic liability in cancer cells^8,9^.

Drug repurposing offers an attractive strategy to accelerate the clinical translation of metabolism-targeting therapies. Clemastine (Clem), a first-generation antihistamine, has emerged as a promyelinating small molecule with sterol-modifying activity and has been linked to inhibition of emopamil binding protein (EBP), a key enzyme in cholesterol biosynthesis^10,11^. Bexarotene (Bex), a selective retinoid X receptor (RXR) agonist, regulates cholesterol transport and efflux pathways through RXR-dependent control of ApoE-, ABCA1-, LDLR-and related lipid-handling programs^12^ . Although these agents have primarily been studied independently in neurological and metabolic disease settings including for their ability to promote oligodendrocyte differentiation and remyelination in demyelinating diseases^10-12^, their potential for affecting cholesterol homeostasis, ER stress adaptation, and lipid stress responses had not previously been explored in GBM^10,12^. Although Bex has shown efficacy in brain diseases, its clinical utility is limited by systemic toxicity. Here, we leverage the unique opportunity for local drug delivery during GBM resection and model this clinically relevant approach in rodents.

Beyond tumor-intrinsic metabolism, GBM progression is strongly shaped by an immunosuppressive, myeloid-rich microenvironment in which cytokine, chemokine, and vascular–stromal signaling influence malignant cell behavior and state transitions^3,7^. Consistent with this framework, our single-cell RNA-seq and intercellular communication analyses indicate that Clem+Bex treatment remodels tumor microenvironment crosstalk, shifting inflammatory and immune-associated signaling networks and redistributing sender–receiver roles across malignant, myeloid, and perivascular/stromal compartments. These findings suggest that this therapy may reprogram not only tumor-intrinsic stress pathways but also the broader immune and vascular ecosystem that supports GBM persistence and recurrence.

Here, we demonstrate that combined Clem and Bex treatment induces profound metabolic disruption in patient-derived glioblastoma models through coordinated perturbation of cholesterol biosynthesis, transport, and storage pathways. This metabolic imbalance is accompanied by intracellular lipid droplet accumulation, sustained ER stress, and activation of all three major UPR branches, ultimately promoting autophagy and apoptotic cell death. Integrated transcriptomic, ultrastructural, and single-cell analyses reveal activation of ER stress, autophagy, and antigen-presentation programs in GBM cells, accompanied by tumor microenvironment remodeling that may enhance antitumor immunity. Notably, Clem+Bex preferentially targets GBM cells, while normal brain cells exhibit minimal stress responses and primarily activate regeneration- and plasticity-associated programs. Together, our findings identify cholesterol dependency as a key metabolic vulnerability in GBM and establish Clem–Bex therapy as a clinically translatable strategy that couples metabolic stress to tumor cell death and immune activation.

## RESULTS

### Clemastine and Bexarotene synergistically suppress proliferation and induce apoptotic cell death in patient-derived GSCs

Previous studies have shown that Clem disrupts cholesterol biosynthesis through inhibition of the late sterol-processing enzyme emopamil binding protein (EBP) although it is not its canonical pathway this has been previously described for glial progenitors ^11,13^, (Fig. S1) whereas Bex modulates cholesterol uptake, transport, and lipid homeostasis through RXR-ABCA1 or RXR-IDOL-LDLR pathways^12-14^. We hypothesized that simultaneous targeting of these complementary pathways would induce metabolic stress and compromise GBM cell survival.

To test this, we first determined the sensitivity of four patient-derived GSC lines (GBM1A, QNS690, QNS712, and QNS986, from our biobank to Clem and Bex individually (Fig. 1A-B, Fig. S2A). Across models, average IC50 values were approximately 15 μM for Clemastine and 35 μM for Bexarotene. Combination screening across multiple dose levels revealed robust synergistic interactions in all GSC lines, with high Highest Single Agent (HSA) synergy scores demonstrating greater-than-additive antitumor activity (Fig. 1C, Fig. S2B). Consistent with these findings, combined treatment produced significantly greater growth inhibition than either agent alone (Fig. 1D, Fig. S2C).

**Figure 1.**
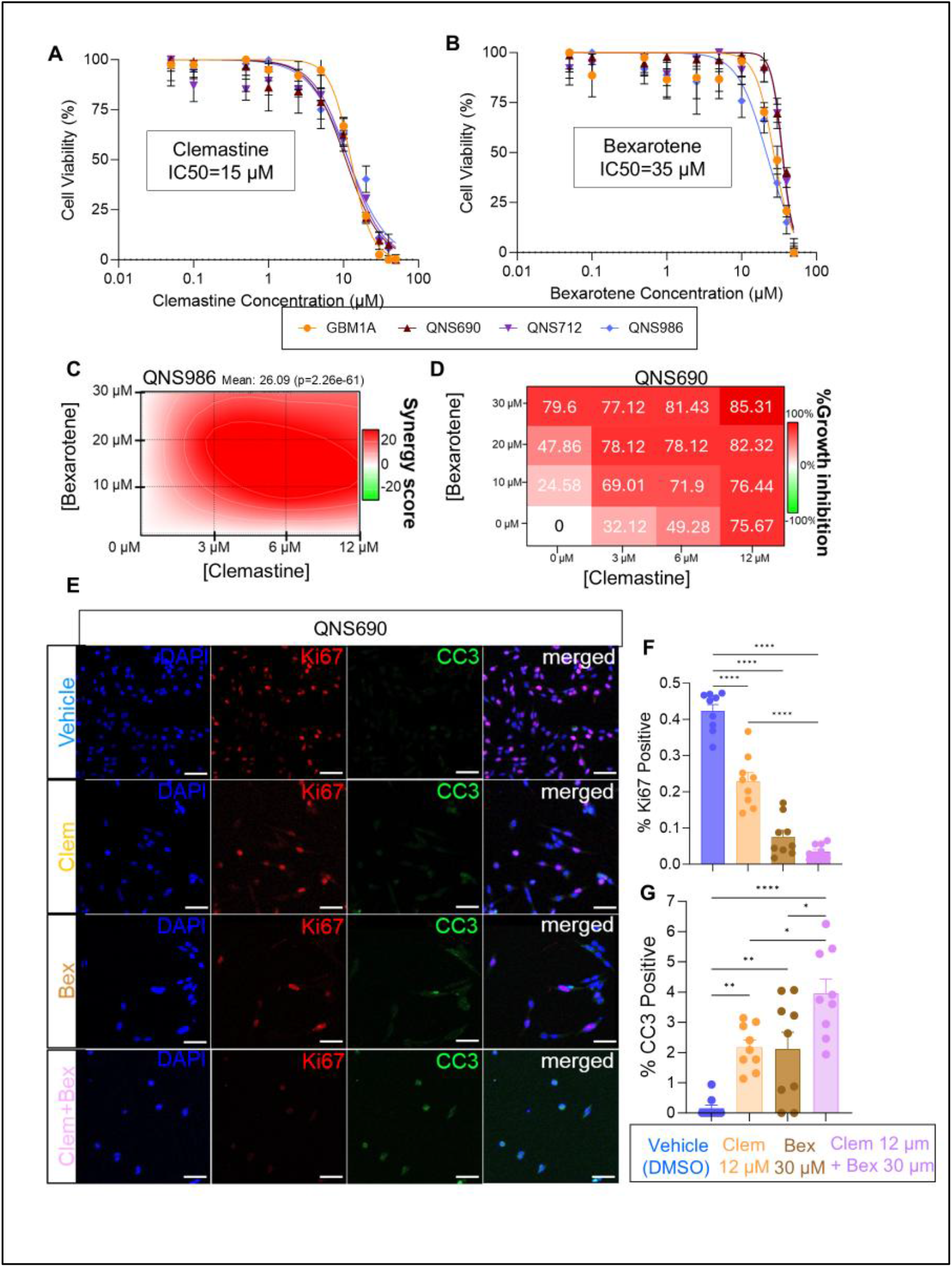
Clemastine and Bexarotene synergistically inhibit growth and induce apoptosis in patient-derived GBM GSCs. A–B) Dose-response curves of Clemastine (A) and Bexarotene (B) in GBM1A, QNS690, QNS712, and QNS986 GSCs; IC₅₀ values were determined by nonlinear regression. C) Synergy analysis of combined Clemastine and Bexarotene in QNS986 cells. D) Heatmap showing growth inhibition in QNS690 cells treated with increasing concentrations of Clemastine and Bexarotene (IC₀, 12.5, 25, and 50). E) Representative immunofluorescence images of Ki67 and cleaved caspase-3 (CC3) in QNS690 cells treated with vehicle, Clem (12 μM), Bex (30 μM), or the combination. Scale bar, 100 μm. F–G) Quantification of Ki67-positive proliferating cells (F) and CC3-positive apoptotic cells (G) across all cell lines. Data are mean ± SEM. One-way ANOVA with Tukey’s multiple-comparisons test. *P < 0.05, **P < 0.01, ***P < 0.001, ****P < 0.0001.

Functional validation confirmed that treatment with either drug reduced GSC proliferation after 72 hours; however, the combination elicited substantially stronger effects across all patient-derived models (Fig. 1E-F, Fig. S3A-B). Similarly, apoptotic cell death was increased by single-agent treatment and was further enhanced following combination therapy (Fig. 1E, G, Fig. S3A, C). Together, these findings demonstrate that Clemastine and Bexarotene synergistically impair GBM cell growth and promote apoptosis across molecularly distinct patient-derived GSC models, supporting the broad therapeutic potential of this metabolic strategy.

### Combined metabolic therapy promotes differentiation and suppresses stem-like programs in GSCs

Both Clem and Bex separately have previously been shown to promote differentiation of oligodendrocyte progenitor cells in demyelinating disorders, including multiple sclerosis ^13,15-17^. Because OPC-like GSCs represent one of the most proliferative and invasive cellular states in GBM3, we next investigated whether the antiproliferative effects of treatment were accompanied by alterations in stemness and differentiation.

Treatment with Clem and Bex for 72 hours significantly reduced expression of the stem cell-associated markers Nestin, Sox2, and Vimentin across all patient-derived GSC lines (Fig. 2A-Fig. S3A, E). While individual agents partially decreased stemness marker expression, the combination consistently produced the strongest effects. In parallel, expression of the glial differentiation marker GFAP was significantly increased following combination treatment (Fig. 2A,C-F, Fig. S3A,D), indicating a shift toward a more differentiated cellular phenotype.

**Figure 2.**
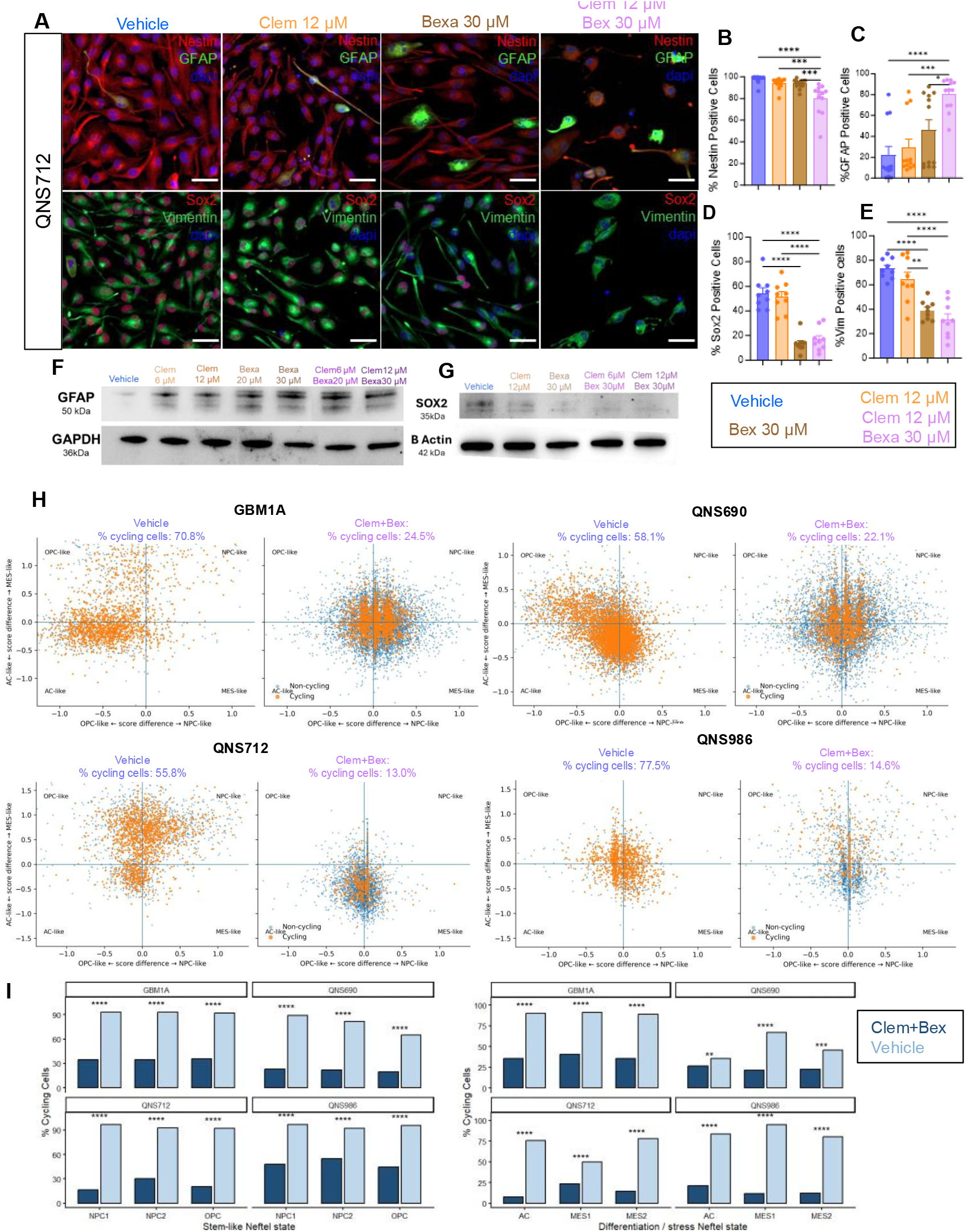
Clemastine and Bexarotene reduce GBM stemness and proliferation while reshaping tumor cell states. A) Representative immunofluorescence images of QNS712 GSCs treated with vehicle, Clem (12 μM), Bex (30 μM), or the combination, stained for Nestin, GFAP, SOX2, and Vimentin. Scale bars, 50 μm. B–E) Quantification of Nestin (B), GFAP (C), SOX2 (D), and Vimentin (E). Combination treatment reduced stemness markers and increased GFAP. Data are mean ± SEM. One-way ANOVA with multiple-comparisons correction. *P < 0.05, **P < 0.01, ***P < 0.001, ****P < 0.0001. F–G) Representative western blots of GFAP (F) and SOX2 (G); GAPDH and β-actin served as loading controls. H) Neftel state analysis of single-cell RNA-seq from GBM1A, QNS690, QNS712, and QNS986 treated with vehicle or Clem+Bex. Butterfly plots show AC-, MES-, NPC-, and OPC-like states with cycling (orange) and non-cycling (blue) cells. Clem+Bex markedly reduced cycling cells across all models. I) Quantification of cycling cells within stem-like (NPC1, NPC2, OPC) and differentiated/mesenchymal-like (AC, MES1, MES2) states, demonstrating broad suppression of proliferative tumor programs. Data are mean ± SEM. Two-sided comparisons between vehicle and Clem+Bex groups. *P < 0.05, **P < 0.01, ***P < 0.001, ****P < 0.0001.

To further characterize the transcriptional consequences of Clem+Bex treatment, we performed scRNA-seq across all four GSC models following vehicle or combination treatment. Analysis of established GBM cellular states, including OPC-like, NPC-like, AC-like, and MES-like programs, revealed a consistent reduction in cycling populations across transcriptional states following treatment (Fig. 2H–I). Across all models, Clem+Bex markedly decreased the overall fraction of cycling cells, reducing cycling populations from 55.8–77.5% under vehicle conditions to 13.0–24.5% following treatment (Fig. 2H, Fig. S4A). Importantly, this reduction was observed not only within stem-like populations, including NPC- and OPC-like states, but also within AC-like and MES-like differentiation/stress-associated states (Fig. 2I). Thus, Clem+Bex broadly suppresses cell-cycle activity across GBM cellular states rather than selectively depleting a single transcriptional compartment. These findings are consistent with reduced proliferative potential and align with the observed decrease in SOX2 expression and increase in GFAP expression following treatment. Collectively, these data suggest that metabolic disruption by Clem+Bex limits GBM growth by broadly suppressing cycling programs across both stem-like and differentiated/stress-associated tumor cell states, while promoting phenotypic changes associated with reduced stemness and increased differentiation.

### Combined Clemastine–Bexarotene therapy disrupts cholesterol homeostasis, leading to lipid droplet accumulation and cholesterol sequestration

Because both Clemastine (Clem) and Bexarotene (Bex) regulate cholesterol metabolism in oligodendrocyte precursor cells, we investigated whether these pathways were similarly altered in patient-derived GSCs. Following 72 h treatment with Clem (12 μM) and Bex (30 μM), bulk RNA-seq and GSEA revealed significant dysregulation of sterol biosynthesis pathways (Fig. S5). Multiple genes involved in cholesterol synthesis, including HMGCR, SQLE, EBP, DHCR24, and DHCR7, were broadly downregulated across all GSC models. These findings were validated at the protein level, where active SREBP1 and SREBP2 were reduced following treatment, indicating suppression of the cholesterol biosynthetic program (Fig. S4B, S5,S6). As previously reported, Clem treatment led to accumulation of EBP, consistent with its direct inhibition of this enzyme. Importantly, combination therapy produced a stronger suppression of cholesterol biosynthesis than either agent alone. Consistent with these findings, lower expression of the cholesterol biosynthetic signature was associated with improved survival in the TCGA-GBM cohort (Fig. S7).

Since GBM cells rely heavily on exogenous cholesterol, we next examined cholesterol transport pathways. Combination therapy increased expression of the cholesterol efflux transporter ABCA1 while reducing APOE and LDLR, and induced the LXR target IDOL, indicating reduced cholesterol uptake and enhanced cholesterol efflux through RXR/LXR signaling (Fig. S6J–N, Fig. S8). These data demonstrate that Clem and Bex cooperatively disrupt cholesterol homeostasis by simultaneously inhibiting cholesterol synthesis, reducing uptake, and promoting efflux.

To determine how these changes affected intracellular cholesterol trafficking, we analyzed genes involved in cholesterol storage and mobilization. GSEA demonstrated enrichment of lipid droplet organization and cholesterol storage pathways following treatment (Fig. 3A–C). Transmission electron microscopy revealed a marked increase in both lipid droplet number and size across all GSC models, while BODIPY-cholesterol staining confirmed accumulation of cholesterol-rich intracellular vesicles (Fig. 3D–H, Fig. S9). Together, these findings indicate that Clem+Bex induces a coordinated collapse of cholesterol homeostasis, preventing cholesterol from being synthesized, imported, or efficiently mobilized, while sequestering sterol intermediates within lipid droplets and depleting the accessible cholesterol pool required for tumor growth (Fig. 3I).

**Figure 3.**
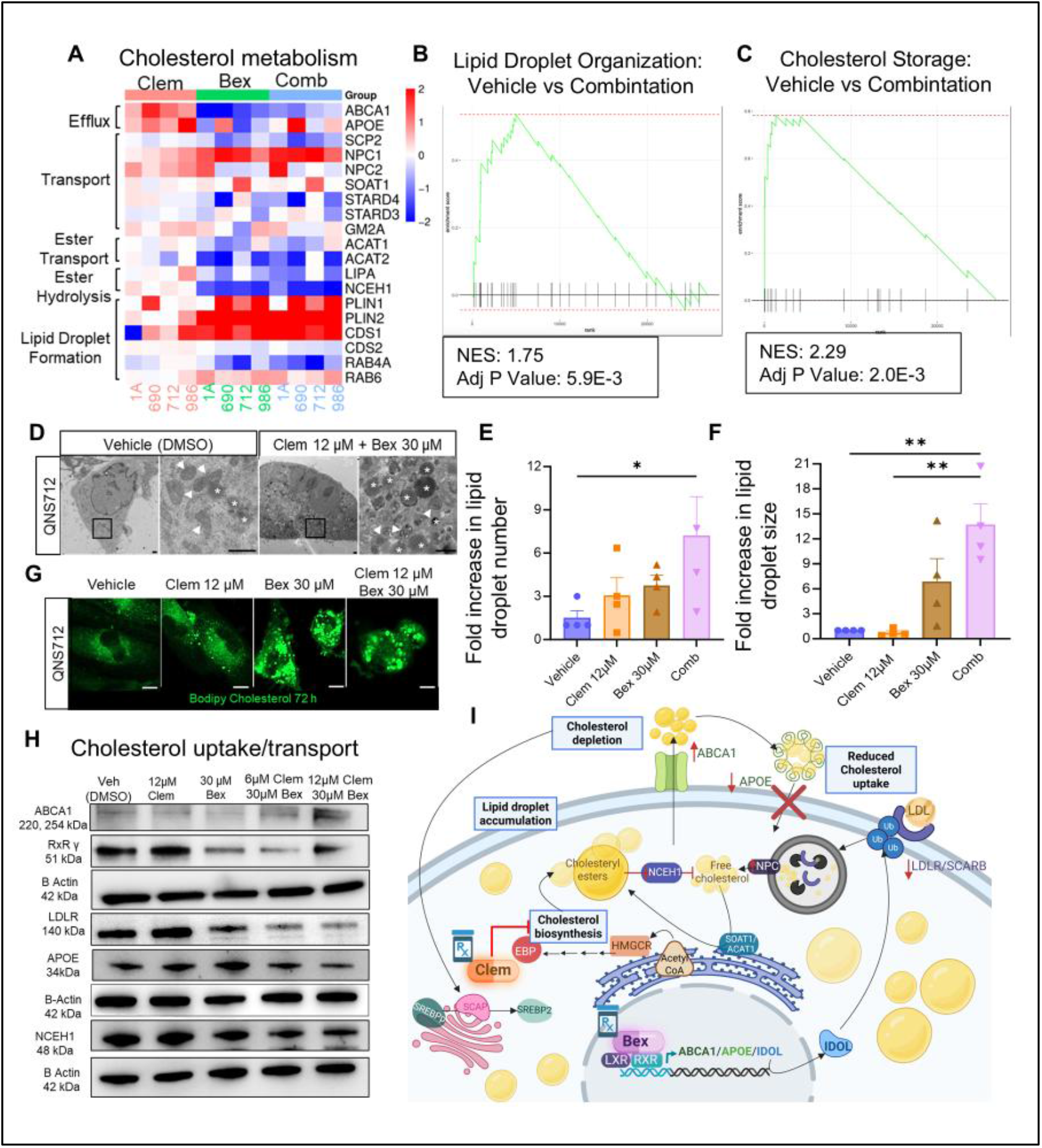
Combined metabolic therapy disrupts cholesterol homeostasis and induces lipid droplet accumulation. A) Heatmap of genes involved in cholesterol metabolism and lipid droplet formation following treatment. B–C) GSEA showing enrichment of lipid droplet organization (B) and cholesterol storage (C) pathways after Clem+Bex treatment. D) Representative TEM images of QNS712 cells treated with vehicle or Clem+Bex (IC₅₀), showing lipid droplet accumulation (white asterisks). Mitochondria are indicated by arrowheads. Scale bars, 1 μm. E–F) Quantification of lipid droplet number (E) and size (F). G) Representative BODIPY cholesterol staining showing intracellular cholesterol accumulation. Scale bars, 5 μm. H) Western blots of ABCA1, RXRγ, LDLR, APOE, and NCEH1, demonstrating reduced cholesterol uptake/transport and increased cholesterol efflux. I) Schematic of Clem+Bex-mediated disruption of cholesterol homeostasis. Data are mean ± SEM. One-way ANOVA with Tukey’s multiple-comparisons test. *P < 0.05, **P < 0.01.

### Cholesterol dysregulation induces endoplasmic reticulum stress and activation of the unfolded protein response

Given the significant role of cholesterol in maintaining membrane composition, protein trafficking, and endoplasmic reticulum (ER) function, we next investigated whether the profound disruption of cholesterol homeostasis induced by Clem and Bex resulted in ER stress. Cholesterol depletion and altered lipid composition have previously been linked to protein misfolding, defective membrane dynamics, and activation of stress response pathways. Consistent with this, pathway enrichment analysis comparing vehicle and combination-treated cells revealed significant upregulation of pathways associated with ER stress, unfolded protein response (UPR), protein processing in the ER, and proteostasis disruption (Fig. 4A). These transcriptional changes were accompanied by marked ultrastructural alterations, including ER dilation, membrane disorganization, and concentric membranous whorl formation, as revealed by TEM across all patient-derived GSC lines (Fig. 4B, Fig. S10A), indicating severe disruption of ER homeostasis.

**Figure 4.**
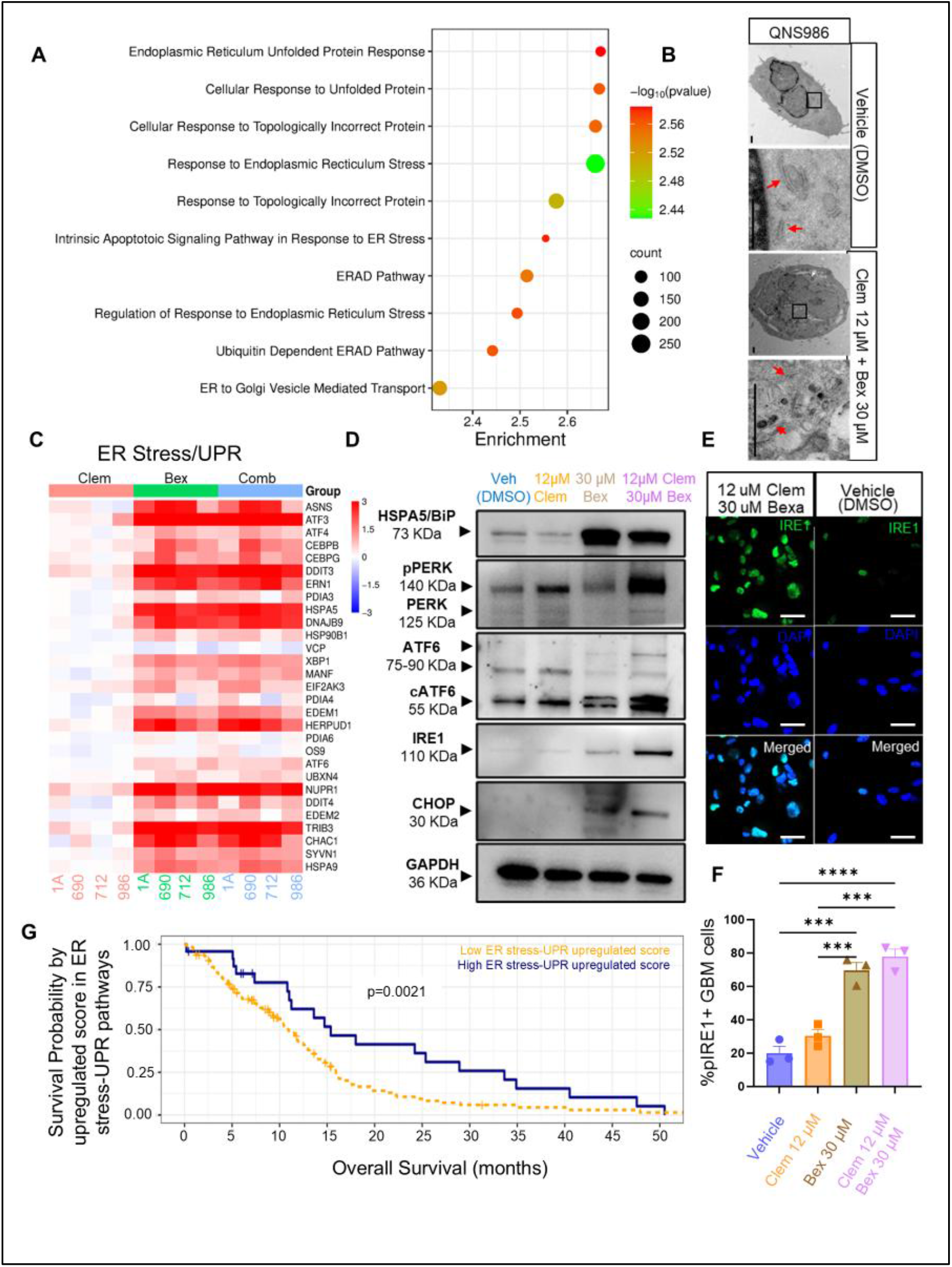
Clemastine+Bexarotene activates ER stress and unfolded protein response pathways. A) Pathway enrichment showing activation of ER stress, UPR, ERAD, and apoptosis following combination therapy. B) Representative TEM images of QNS986 cells showing ER whorls (red arrows). Scale bars, 1 μm. C) Heatmap of ER stress- and UPR-related genes. D) Western blots of BiP, pPERK, PERK, ATF6, cleaved ATF6, IRE1, and CHOP. E–F) Representative IRE1 immunofluorescence (E) and quantification of pIRE1-positive cells (F). Scale bars, 25 μm. G) Kaplan–Meier analysis of the TCGA-GBM cohort showing improved survival in patients with high ER stress-UPR signature scores. Data are mean ± SEM. One-way ANOVA with Tukey’s multiple-comparisons test.*P < 0.05, **P < 0.01, ***P < 0.001, ****P < 0.0001.

We next examined the expression of key regulators of the ER stress response. Combination treatment resulted in robust activation of the three canonical UPR signaling branches. Specifically, expression of the ER stress sensors EIF2AK3 (PERK), ATF6, and ERN1 (IRE1) was significantly increased, together with the master ER chaperone HSPA5 (BiP/GRP78), which functions as the primary regulator of UPR activation (Fig. 4C–F, Fig. S10 B-C). Activation of these pathways was accompanied by increased expression of downstream mediators involved in protein quality control, autophagy, and apoptotic signaling. Notably, the pro-apoptotic transcription factor DDIT3 (CHOP) was strongly induced following treatment, indicating progression beyond an adaptive stress response toward terminal UPR signaling.

Under physiological conditions, activation of the UPR serves as a protective mechanism that restores ER homeostasis by reducing protein synthesis and enhancing protein folding capacity. However, when ER stress is prolonged or excessive, these compensatory mechanisms become insufficient and the UPR transitions into a cell death program ^9,26^. The simultaneous induction of ER stress sensors, autophagy-related pathways, and apoptotic mediators observed in our datasets suggests that Clem and Bex drive GBM cells toward an irreversible ER stress state characterized by persistent UPR activation and commitment to cell death.

To assess the clinical relevance of this response, we generated an ER stress/UPR gene signature based on genes significantly induced by combination treatment and interrogated the TCGA-GBM cohort. Kaplan–Meier survival analysis demonstrated that patients with elevated expression of this signature exhibited significantly prolonged overall survival compared with patients with low expression (Fig. 4G). These findings indicate that activation of ER stress and terminal UPR programs is associated with improved clinical outcome and support ER stress induction as a key mechanism underlying the antitumor activity of combined Clem and Bex treatment.

### Irreversible ER stress promotes autophagy-mediated cell death

Because sustained activation of the three major UPR branches (ATF6, IRE1, and PERK) is known to trigger autophagy and apoptosis when ER homeostasis cannot be restored9,46, we next investigated whether the terminal UPR state induced by Clem and Bex resulted in activation of autophagic cell death pathways. Pathway enrichment analysis revealed significant upregulation of autophagy-related programs, which was further confirmed by GSEA demonstrating enrichment of pathways associated with positive regulation of autophagy (Fig. 5A).

**Figure 5.**
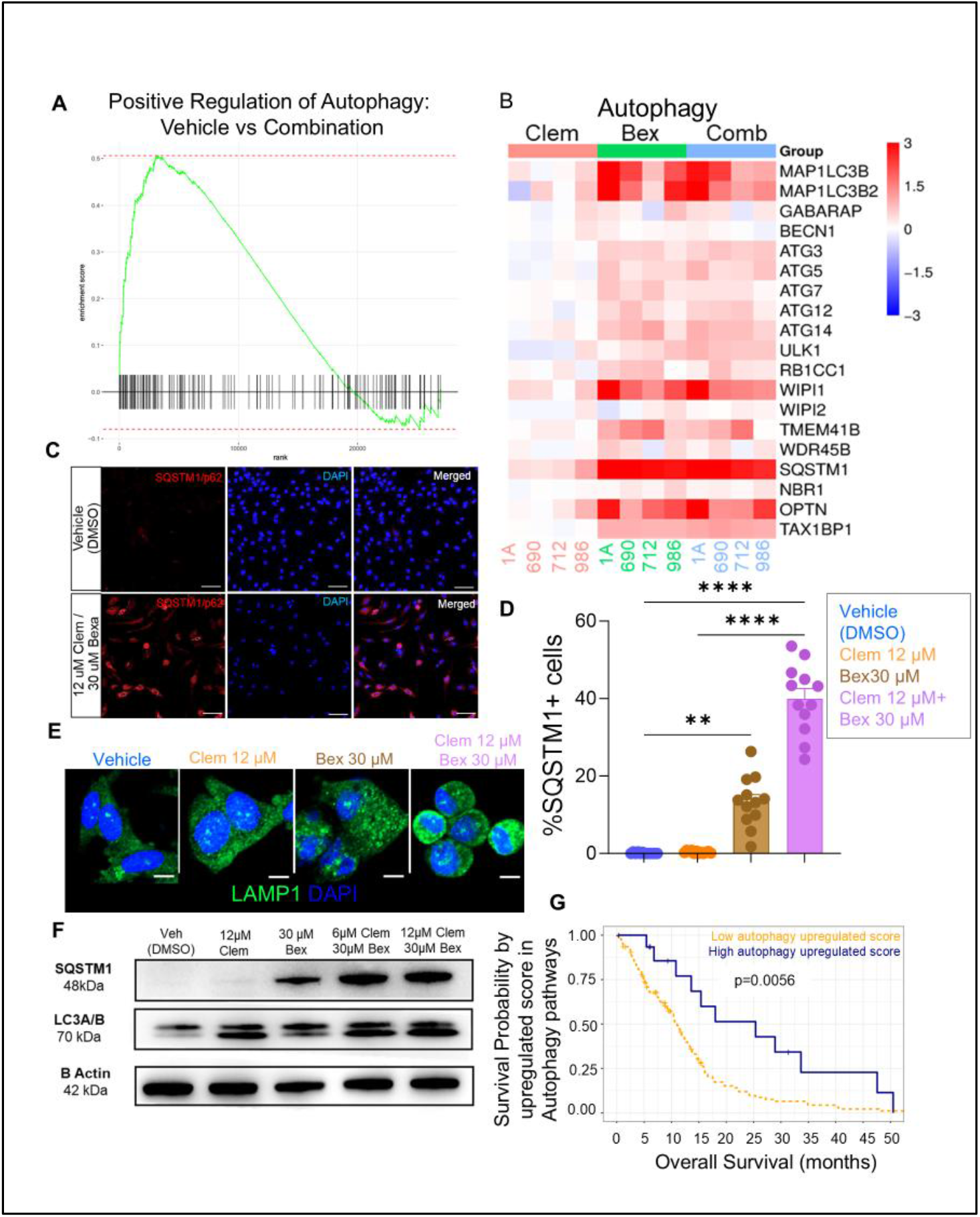
Clemastine+Bexarotene induces autophagy signaling. A) GSEA showing enrichment of autophagy pathways following combination therapy. B) Heatmap of autophagy-associated genes across all four patient-derived GSC models. C–D) Representative SQSTM1/p62 immunofluorescence images (C) and quantification (D) showing increased p62 expression after treatment. Scale bars, 50 μm. E) Representative LAMP1 immunofluorescence demonstrating increased lysosomal abundance. Scale bars, 20 μm. F) Western blots of SQSTM1/p62 and LC3A/B. G) Kaplan–Meier analysis of the TCGA-GBM cohort showing improved survival in patients with high autophagy signature scores. Data are mean ± SEM. One-way ANOVA with Tukey’s multiple-comparisons test. *P < 0.05, **P < 0.01, ***P < 0.001, ****P < 0.0001.

At the transcriptional level, combination treatment induced robust expression of key autophagy regulators, including MAP1LC3B (LC3B) and SQSTM1/p62, two central mediators of autophagosome formation and cargo degradation. We additionally observed upregulation of WIPI1, a critical effector that links phospholipid signaling to phagophore initiation through recruitment of autophagy-related proteins (ATGs), indicating active autophagosome biogenesis (Fig. 5B–E, Fig. S11). Notably, OPTN (Optineurin) was also significantly increased following treatment. Because OPTN functions as a selective autophagy receptor responsible for targeting damaged organelles and misfolded protein aggregates for degradation, its induction provides a direct mechanistic link between persistent ER stress, proteotoxic damage, and activation of selective autophagy pathways.

To further validate autophagic flux, we examined expression of LAMP1, a lysosomal membrane protein required for autophagosome maturation and lysosomal fusion. LAMP1 was significantly upregulated following treatment, supporting enhanced autophagosome processing and lysosome-dependent degradation (Fig. 5F). Together, these findings indicate that cholesterol deprivation-induced ER stress progresses beyond adaptive UPR signaling and activates a robust autophagic program aimed at eliminating damaged cellular components. However, because this response occurs concomitantly with activation of terminal UPR pathways and pro-apoptotic mediators, our data suggest that autophagy ultimately contributes to cell death rather than cellular recovery.

To evaluate the clinical relevance of these findings, we generated an autophagy-associated gene signature using treatment-induced genes and interrogated the TCGA-GBM cohort. Kaplan–Meier analysis demonstrated that elevated expression of this signature was associated with significantly prolonged overall survival, indicating that activation of autophagy-related pathways correlates with improved patient outcomes and supporting autophagy-mediated cell death as a major downstream consequence of combined Clem and Bex treatment (Fig. 5G).

### Intracranial delivery of clemastine and bexarotene enhances therapeutic efficacy in orthotopic GBM through local target engagement and improved drug exposure

Given that both Clem and Bex have been evaluated in neurological disorders, including multiple sclerosis (MS), their pharmacokinetic properties are well characterized. Clem readily crosses the gastrointestinal tract and blood–brain barrier (BBB), achieving therapeutic CNS concentrations^15,17^. In contrast, despite its lipophilic nature, only ∼13% of systemically administered Bex reaches the CNS12,27,28, consistent with in silico BBB permeability predictions (Fig. 6A). These findings suggest that limited intratumoral drug exposure, particularly for Bex, may reduce the efficacy of systemic administration.

**Figure 6.**
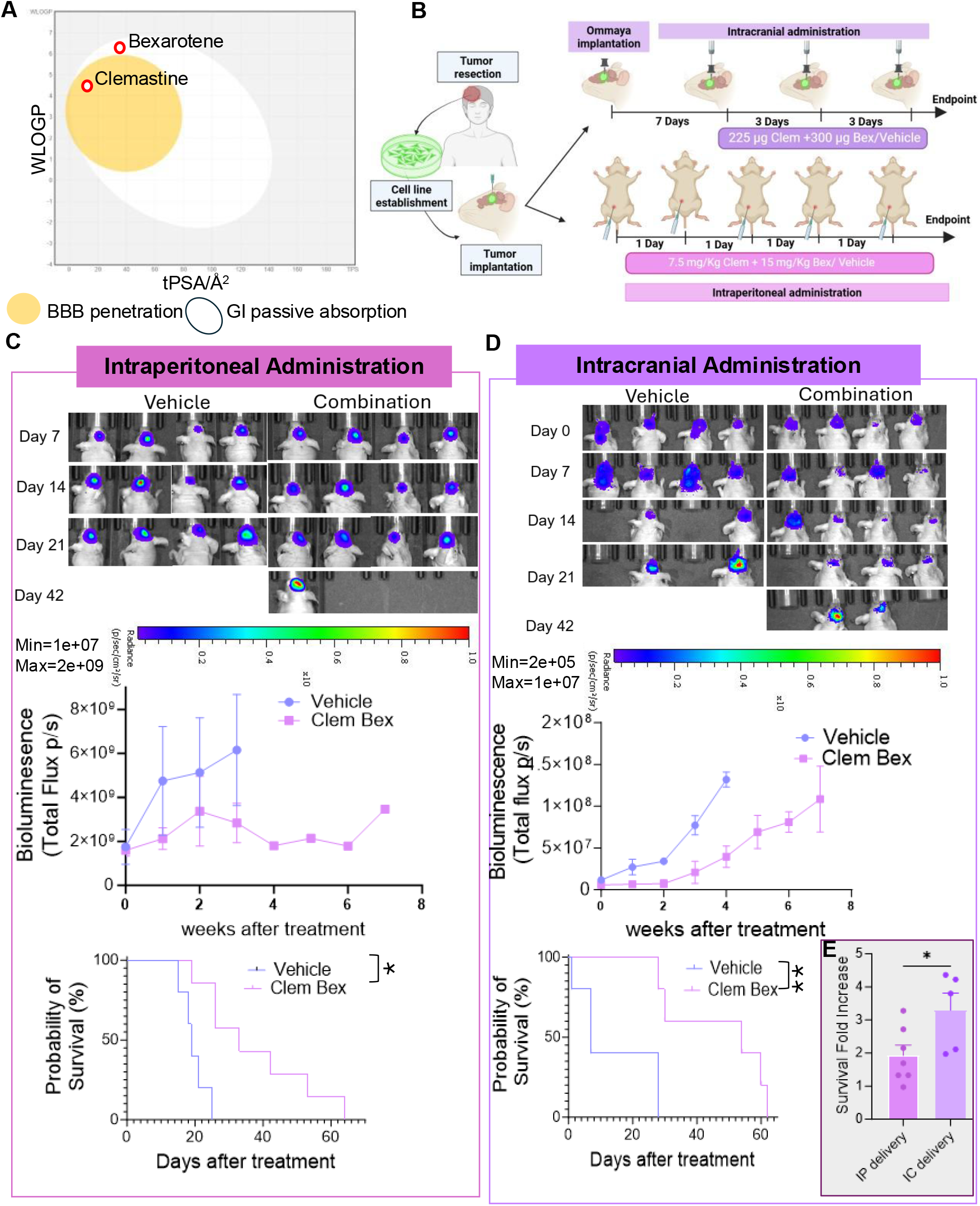
Intracranial delivery of Clemastine+Bexarotene improves efficacy at lower doses in orthotopic GBM models. A) Schematic of blood–brain barrier penetration and gastrointestinal absorption of Clemastine and Bexarotene, highlighting the limited BBB penetration of Bexarotene. B) Experimental design comparing intraperitoneal and intracranial administration. C–D) Representative bioluminescence imaging, tumor growth, and Kaplan–Meier survival analyses following intraperitoneal (C) or intracranial (D) treatment. E) Survival benefit normalized to vehicle, demonstrating superior efficacy of intracranial delivery at lower doses. Data are mean ± SEM. *P < 0.05, **P < 0.01

Because systemic Bex is also associated with dose-limiting toxicities^29-31^, we hypothesized that direct intracranial delivery could maximize tumor exposure while minimizing systemic toxicity. We therefore compared systemic and intracranial Clem+Bex administration in orthotopic GBM models. Mice received either intraperitoneal Clem (7.5 mg/kg) and Bex (15 mg/kg) five days per week^19,27,32-34^ or intracranial treatment through an implanted Ommaya reservoir every three days using approximately two-fold lower Clem and four-fold lower Bex doses (Fig. 6B).

Longitudinal bioluminescence imaging demonstrated that both delivery routes significantly reduced tumor growth compared with vehicle controls (Fig. 6C–D). However, intracranial treatment produced a substantially greater survival benefit despite the markedly lower cumulative drug doses. When survival was normalized to the corresponding vehicle controls, intracranial delivery significantly outperformed systemic administration, highlighting the importance of bypassing the BBB and maintaining sustained intratumoral drug concentrations (Fig. 6E). Together, these findings demonstrate that local Clem+Bex administration improves therapeutic efficacy while reducing overall drug exposure, supporting intracranial delivery as a clinically translatable strategy for targeting GBM metabolism.

### In vivo orthotopic models recapitulate the stress–ER stress–autophagy axis observed in vitro

To determine whether the mechanisms identified in vitro were recapitulated in vivo, we established orthotopic GBM models using GBM1A and QNS690 patient-derived GSCs. Following tumor engraftment, mice received intracranial Clem+Bex through an implanted Ommaya reservoir and were sacrificed 72 h later to match the in vitro treatment window. H&E and human nuclei (HuNu) staining confirmed tumor establishment in all mice (Fig. 7A, Fig. S12).

**Figure 7.**
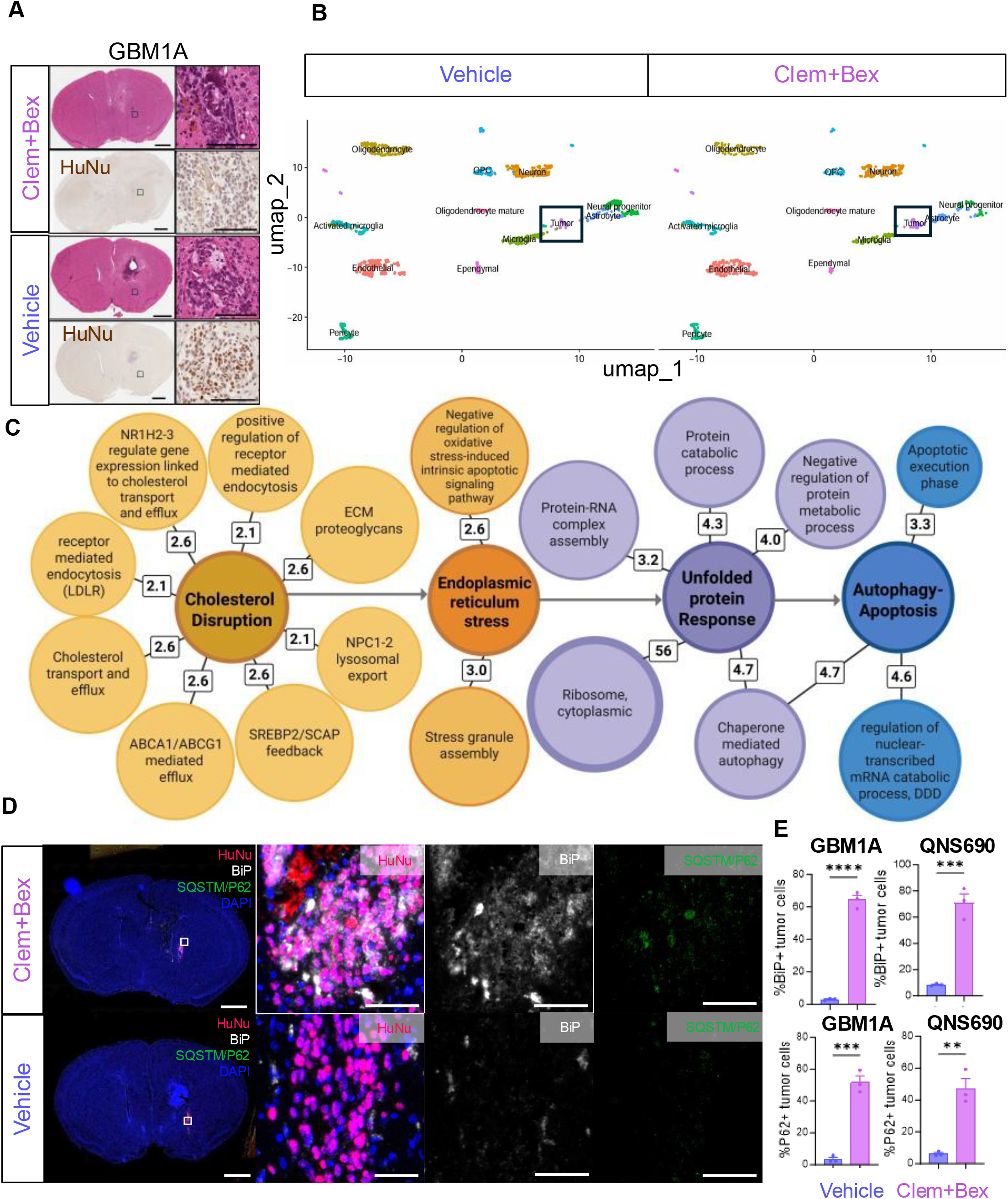
Clemastine+Bexarotene induces cholesterol dysregulation, ER stress, UPR, and autophagy in orthotopic GBM. A) Representative H&E and HuNu staining of orthotopic GBM1A tumors 72 h after intracranial vehicle or Clem+Bex treatment. Scale bars, 1 mm and 25 μm. B) UMAP of single-cell RNA-seq identifying major brain and tumor cell populations, showing no changes in normal brain cell composition following treatment. C) Network analysis linking cholesterol dysregulation with ER stress, UPR, and autophagy/apoptosis pathways. D) Representative BiP and SQSTM1/p62 immunofluorescence in orthotopic tumors, showing increased ER stress and autophagy after treatment. HuNu identifies tumor cells. Scale bars, 1 mm and 50 μm. E) Quantification of BiP- and SQSTM1/p62-positive tumor cells in GBM1A and QNS690 models. Data are mean ± SEM. Unpaired two-tailed Student’s t test. **P < 0.01, ***P < 0.001, ****P < 0.0001.

Single-cell RNA sequencing of GBM1A tumors identified 12 cellular populations, including 11 murine brain cell types and one human tumor cluster (cluster 9) (Fig. S13). Because our goal was to define tumor-specific responses, subsequent analyses focused on the human tumor compartment. UMAP analysis showed no major changes in overall brain cell composition following treatment, indicating that Clem+Bex primarily induced transcriptional rather than cellular remodeling (Fig. 7B, Fig. S13A).

Differential expression and pathway analyses of tumor cells demonstrated activation of the same pathways observed in vitro, including cholesterol dysregulation, ER stress, unfolded protein response, autophagy, and apoptosis (Fig. 7C, Fig. S14A). In contrast, surrounding neurons, astrocytes, and oligodendrocytes showed minimal ER stress or cell death signatures, instead exhibiting pathways associated with cellular maintenance and repair, suggesting a tumor-selective response (Fig. S14B–D).

Multiplex immunofluorescence in both GBM1A and QNS690 tumors confirmed these findings, showing significantly increased BiP/HSPA5 and SQSTM1/p62 expression following treatment (Fig. 7D–E, Fig. S12B). Together, these results demonstrate that intracranial Clem+Bex recapitulates in vivo the mechanistic cascade observed in vitro, characterized by disruption of cholesterol homeostasis, activation of ER stress and the UPR, induction of autophagy, and tumor cell death.

### Low-dose intracranial Clemastine+ Bexarotene therapy remodels the tumor immune microenvironment without inducing systemic lymphopenia

Because systemic Bex has been associated with lymphopenia and immune suppression^19-32,35^, we evaluated whether our low-dose intracranial regimen altered peripheral immunity while simultaneously assessing its effects on the tumor immune microenvironment. CT-2A tumors were established in immunocompetent C57BL/6 mice, followed by intracranial Clem+Bex treatment through an implanted Ommaya reservoir. Mice were sacrificed 72 h after treatment to capture the early immune responses.

Importantly, intracranial treatment did not alter peripheral immune populations or induce systemic lymphopenia (Fig. S15A). Within the tumor microenvironment, most immune populations remained unchanged, although CD4⁺ T-cell infiltration significantly increased following treatment (Fig. 8A, Fig. S15B), suggesting local immune activation rather than systemic immunosuppression.

**Figure 8.**
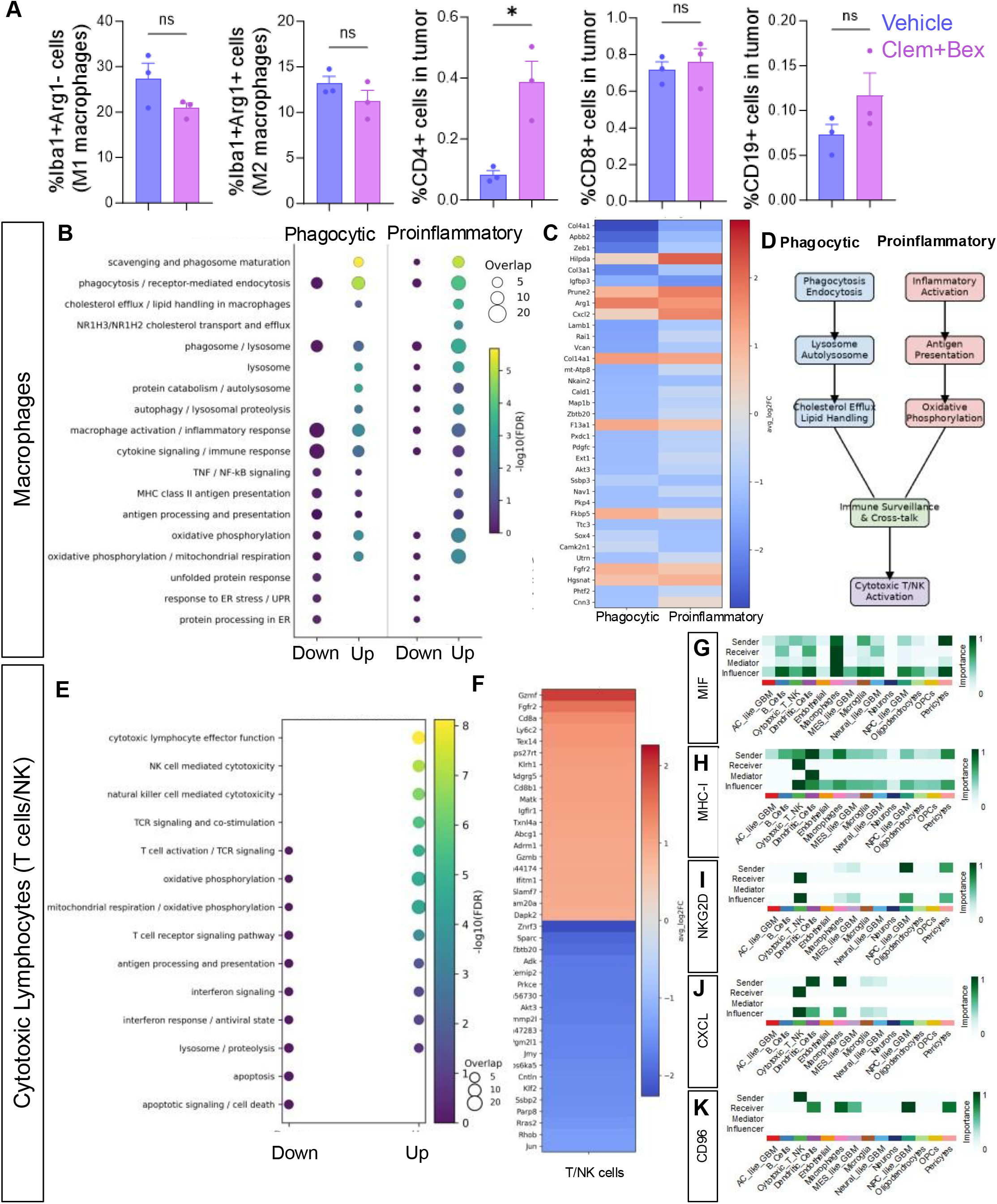
Clemastine+Bexarotene remodels the immune microenvironment and enhances immune communication. A) Quantification of immune cell populations by mIHC in orthotopic syngeneic GBM, showing increased CD4⁺ T cells while preserving macrophages, CD8⁺ T cells, and B cells. Data are mean ± SEM. *P < 0.05; ns, not significant. B) Pathway enrichment of macrophage differentially expressed genes following treatment. C) Heatmap of representative macrophage genes. D) Schematic of Clem+Bex-induced macrophage remodeling, highlighting enhanced phagocytosis, antigen presentation, oxidative metabolism, ER stress, and immune surveillance. E) Pathway enrichment of cytotoxic lymphocyte (T/NK) genes. F) Heatmap of representative cytotoxic lymphocyte genes. G–K) CellChat analysis of MIF (G), MHC-I (H), NKG2D (I), CXCL (J), and CD96 (K) signaling, demonstrating enhanced antigen presentation, chemokine signaling, and cytotoxic immune communication following treatment.

Single-cell RNA sequencing identified major brain-resident cell types, immune populations, and four GBM transcriptional states (AC-, NPC-, MES-, and Neu-like) (Figs. S16–S17). Consistent with our human orthotopic and in vitro datasets, all tumor states exhibited activation of cholesterol dysregulation, ER stress, UPR, and proteotoxic stress pathways, together with enrichment of antigen presentation, inflammatory signaling, cytokine responses, and innate immune activation (Fig. S17B), indicating increased tumor immunogenicity following treatment.

Macrophage analyses revealed two complementary responses. Activated macrophages were enriched for phagocytosis, lipid uptake, and cholesterol processing pathways, consistent with clearance of dying tumor cells, whereas pro-inflammatory macrophages upregulated oxidative phosphorylation, antigen presentation, interferon signaling, and inflammatory programs, indicative of enhanced immune activation (Fig. 8B–D). Cytotoxic lymphocytes similarly exhibited increased expression of genes associated with cytotoxicity and effector function, accompanied by reduced apoptotic programs (Fig. 8E–F).

Finally, CellChat analysis demonstrated that although overall intercellular communication decreased, immune-specific signaling through MIF, MHC-I, NKG2D, CXCL, and CD96 was selectively enhanced (Fig. 8G–K, Fig. S18). Together, these findings indicate that intracranial Clem+Bex not only induces tumor cell death through cholesterol dysregulation, ER stress, and autophagy, but also remodels the tumor immune microenvironment by promoting phagocytosis, antigen presentation, and cytotoxic immune surveillance.

## DISCUSSION

GBM remains one of the most treatment-resistant cancers, despite standard of care which includes maximal surgical safe resection, followed by chemoradiation, in part because of its extraordinary ability to adapt to metabolic stress^36-38^. Here, we show that combined Clem and Bex treatment exploits a previously underexplored vulnerability in GBM by disrupting cholesterol homeostasis. Across multiple patient-derived GSC models, the combination reduced proliferation, decreased stemness, increased apoptosis, and prolonged survival in orthotopic tumors. Mechanistically, these effects were accompanied by profound alterations in cholesterol processing, accumulation of lipid droplets, activation of ER stress and UPR pathways, induction of autophagy-associated programs, and ultimately tumor cell death. Together, our findings identify cholesterol-processing dysfunction as a mechanism capable of overriding the adaptive capacity of GSCs.

Our data suggest that the primary consequence of treatment is not cholesterol depletion but failure of cholesterol utilization. This distinction is particularly relevant in GBM, which develops within one of the most cholesterol-rich environments in the body. Astrocytes continuously supply ApoE-containing lipoproteins^23,39-41^, while macrophages and microglia recycle cholesterol released during myelin destruction during tumor invasion^21,22,42^. Rather than relying exclusively on de novo cholesterol synthesis, GBM cells are therefore positioned to exploit abundant extracellular sterol sources^24,43^. The striking lipid droplet accumulation observed across all models argues that cholesterol remains available but cannot be efficiently processed. Instead, excess sterols appear to be diverted into storage compartments, creating a state in which cholesterol is abundant yet functionally inaccessible. The coordinated changes observed in cholesterol uptake, transport, efflux, and storage pathways further support this interpretation. We hypothesize that this disconnection between sterol availability and sterol utilization represents a major source of metabolic stress in treated tumors.

The ER is particularly sensitive to disruptions in cholesterol homeostasis because it coordinates both sterol sensing and proteostasis^9,44,45^. Consistent with this, combined treatment activated all three major branches of the unfolded protein response, including PERK, IRE1, and ATF6 signaling^9,46^. The induction of BiP, CHOP, and other ER stress-associated pathways suggests that GBM cells initially attempt to adapt to sterol-processing defects through activation of protective stress responses^9,46^. However, these compensatory mechanisms appear insufficient to restore homeostasis. Instead, persistent lipid imbalance drives chronic ER stress that progressively shifts from an adaptive to a pro-death response. This interpretation is supported by the robust induction of autophagy-associated pathways observed across transcriptomic, protein, and imaging datasets. Autophagy likely functions initially as an attempt to recycle damaged cellular components and mobilize intracellular nutrient stores. Yet despite this response, tumor cells ultimately undergo apoptosis, indicating that the magnitude of metabolic stress exceeds their capacity for adaptation. Together, these findings place ER stress and UPR activation at the center of the biological response to cholesterol dysregulation in GBM.

The single-cell analyses suggest that the effects of metabolic therapy extend beyond direct tumor cell killing. In addition to reducing proliferative and stem-like tumor states, treatment induced transcriptional programs associated with regeneration, tissue remodeling, and cellular plasticity within the neural and glial tumor microenvironment. Furthermore, metabolic therapy profoundly reshaped the immune tumor microenvironment, promoting activation of inflammatory and antigen-presentation pathways, expansion of phagocytic and pro-inflammatory macrophage states, and enhanced cytotoxic lymphocyte programs. These changes are consistent with a transition from an immunologically "cold" tumor microenvironment toward a more immune-responsive state capable of supporting antitumor immunity. Together, these findings indicate that metabolic stress not only suppresses tumor growth but also reprograms both resident brain cells and infiltrating immune populations. Notably, activation of regeneration and plasticity pathways may reflect an adaptive response to treatment-induced injury, whereas activation of innate and adaptive immune programs likely contributes to the clearance of stressed and dying tumor cells. Understanding these coordinated responses may uncover additional opportunities to enhance therapeutic efficacy, promote durable tumor control, and rationally combine metabolic therapy with immunotherapeutic approaches.

Some limitations of the present study should be acknowledged. While our data establish cholesterol dysregulation, ER stress, and autophagy as central features of the therapeutic response, the relative contributions of cholesterol uptake, trafficking, storage, and biosynthesis remain to be fully defined. Furthermore, the functional significance of the regeneration and plasticity programs identified by single-cell analyses will require additional investigation in models that closely recapitulate the human disease or in the human brain directly in the future. Despite these limitations, our findings provide a mechanistic framework linking cholesterol homeostasis to GBM survival and identify metabolic disruption as a promising therapeutic strategy for this disease.

The most encouraging aspect of this study is its translational potential. Both Clem and Bex are clinically available compounds, yet intracranial administration achieved substantial antitumor activity using doses far below those typically required systemically. This finding underscores the importance of local drug delivery for metabolic therapies targeting brain tumors. Importantly, we did not observe major alterations in brain or immune cell composition in orthotopic or syngeneic models, despite previous reports of systemic Bex-associated hematologic toxicity using local delivery. Although additional studies will be required to define the relative contributions of cholesterol uptake, trafficking, storage, and biosynthesis to the observed phenotype, the overall biological framework is clear. Combined Clem and Bex treatment disrupt cholesterol handling, leading to accumulation of unusable sterol intermediates, lipid droplet expansion, ER stress, autophagy-associated remodeling, and apoptotic cell death. These findings establish cholesterol-processing dysfunction as a therapeutically exploitable vulnerability in GBM and provide a rationale for further development of metabolism-based therapeutic strategies for this disease.

### Limitations of the study

This study has several limitations. First, although we demonstrate that coordinated disruption of cholesterol homeostasis induces tumor-intrinsic stress and promotes antitumor immune responses, the contribution of specific cholesterol intermediates and lipid species to these effects remains to be fully defined. Second, while therapeutic activity was validated across multiple patient-derived GBM models and in vivo systems, these models cannot fully recapitulate the cellular and immune complexity of human GBM. Finally, although intracranial administration enabled substantial dose reduction and effective local treatment, optimal dosing, distribution, and delivery parameters will require further investigation before clinical translation.

## STAR METHODS

### KEY RESOURCES TABLE

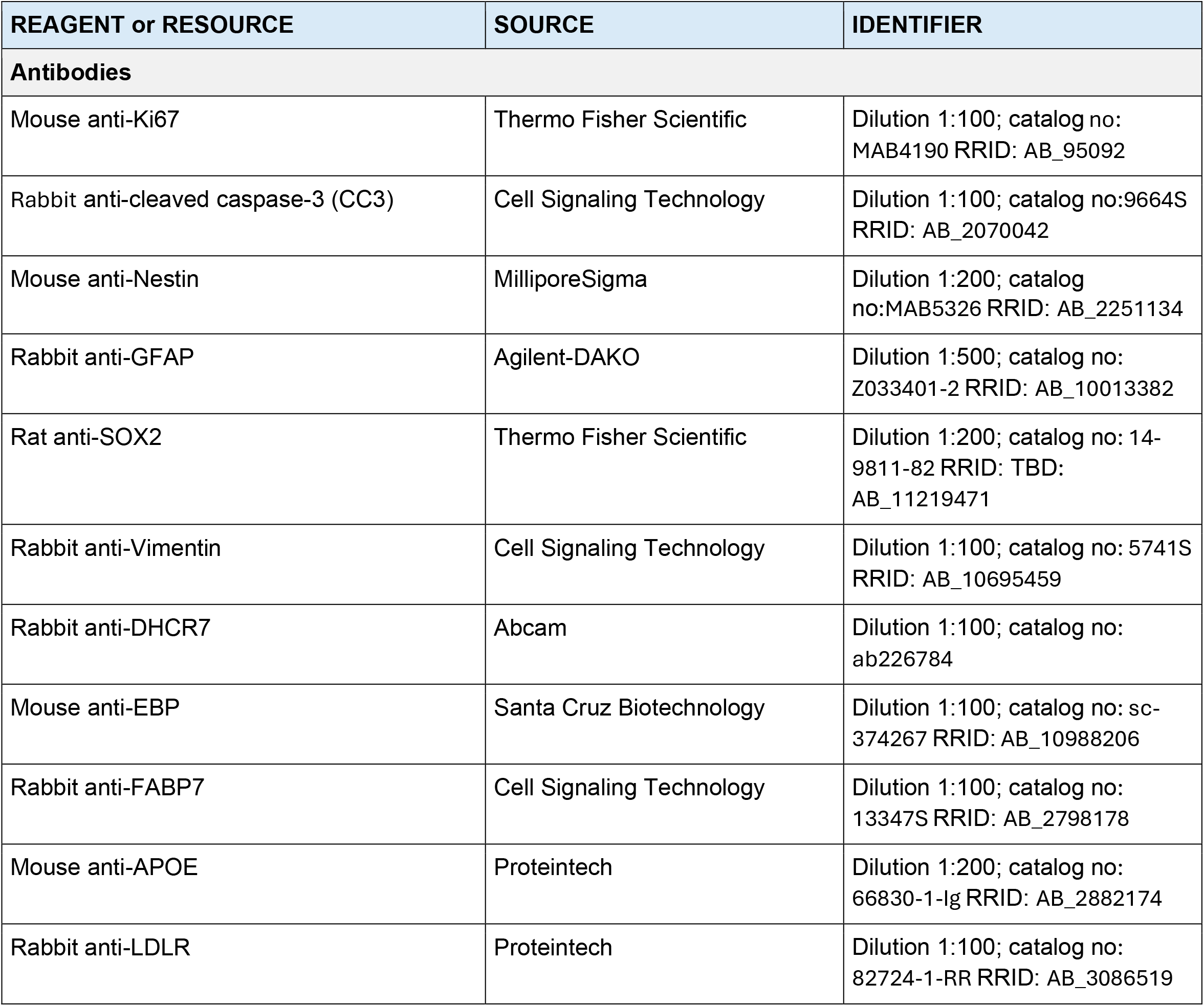
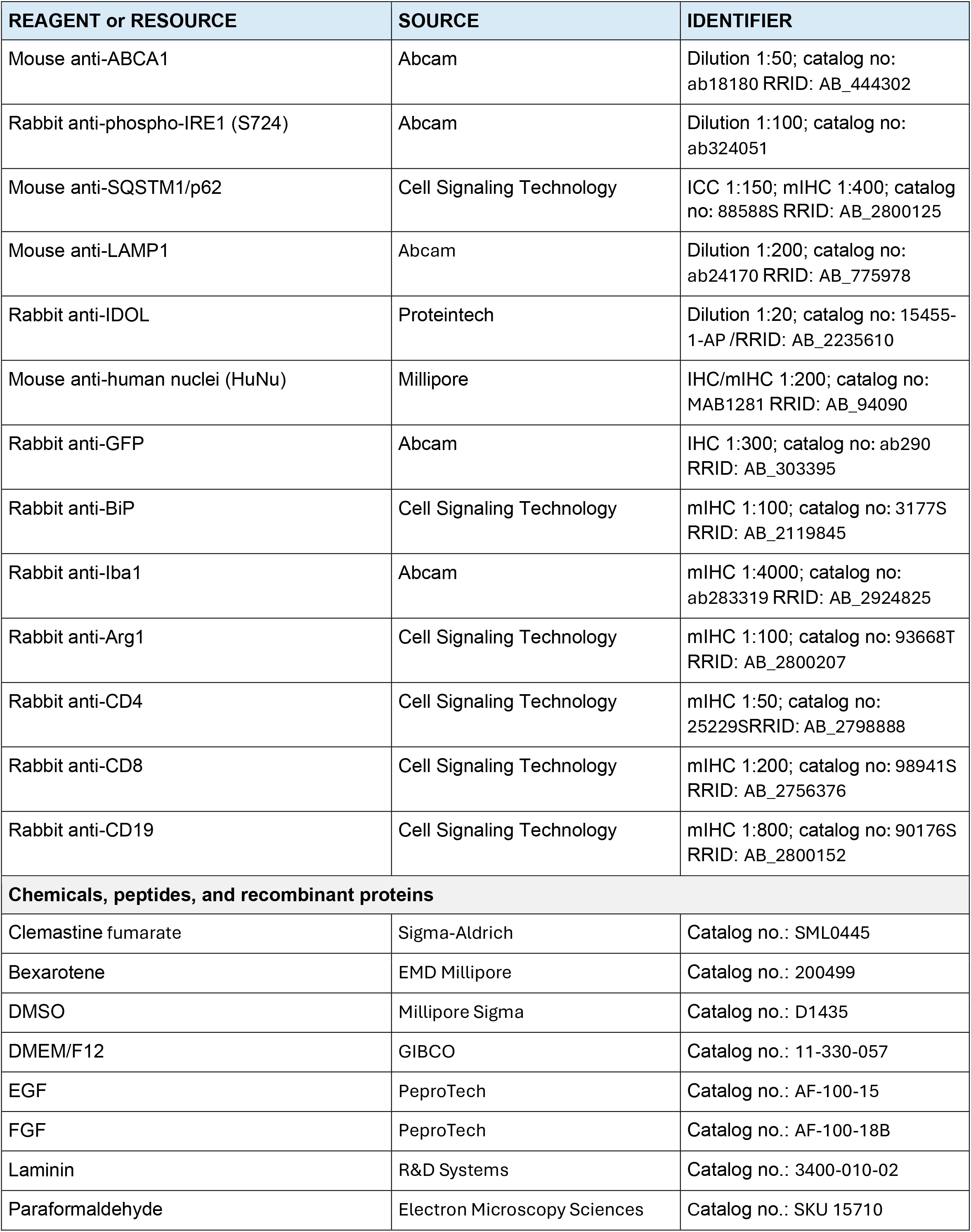
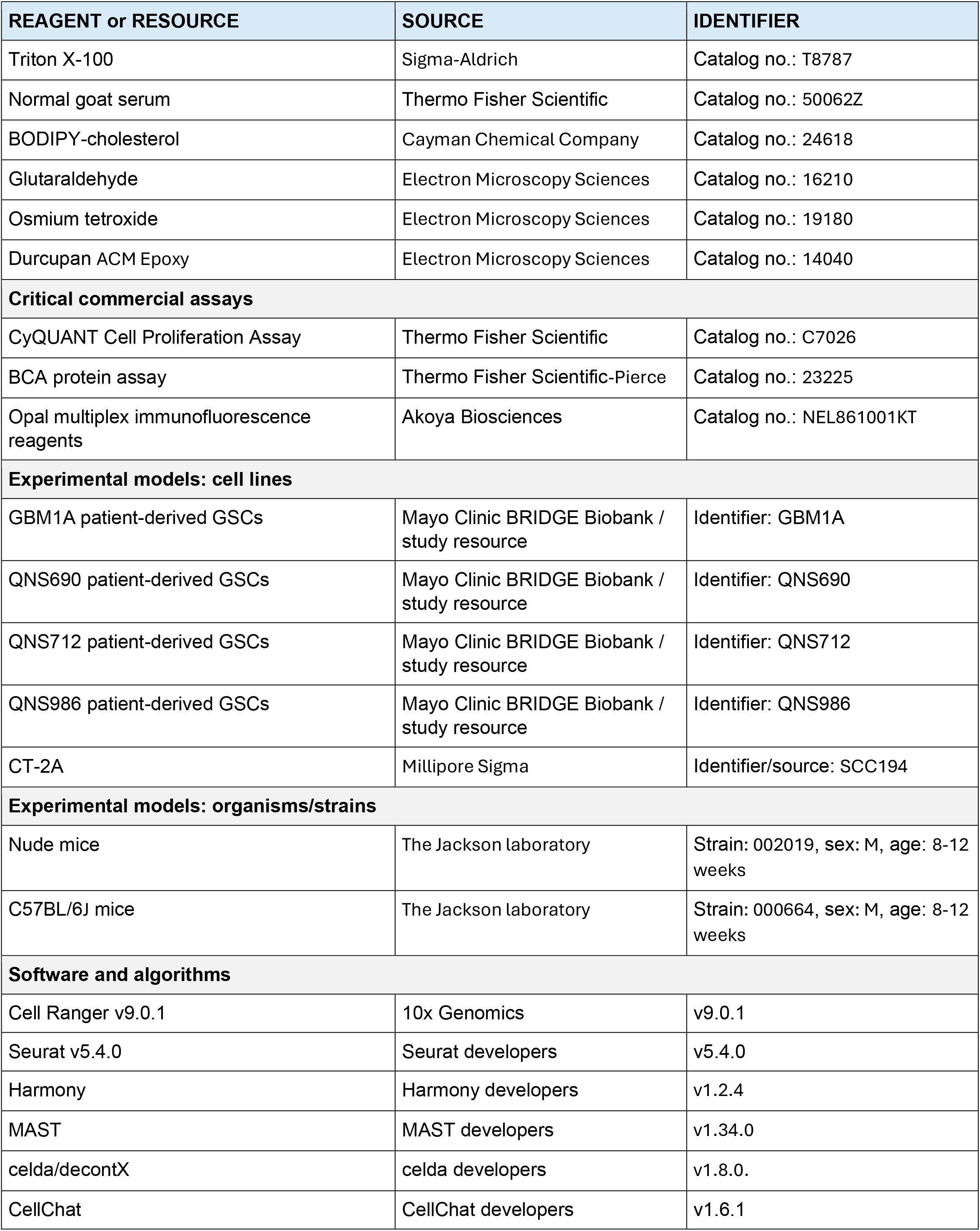
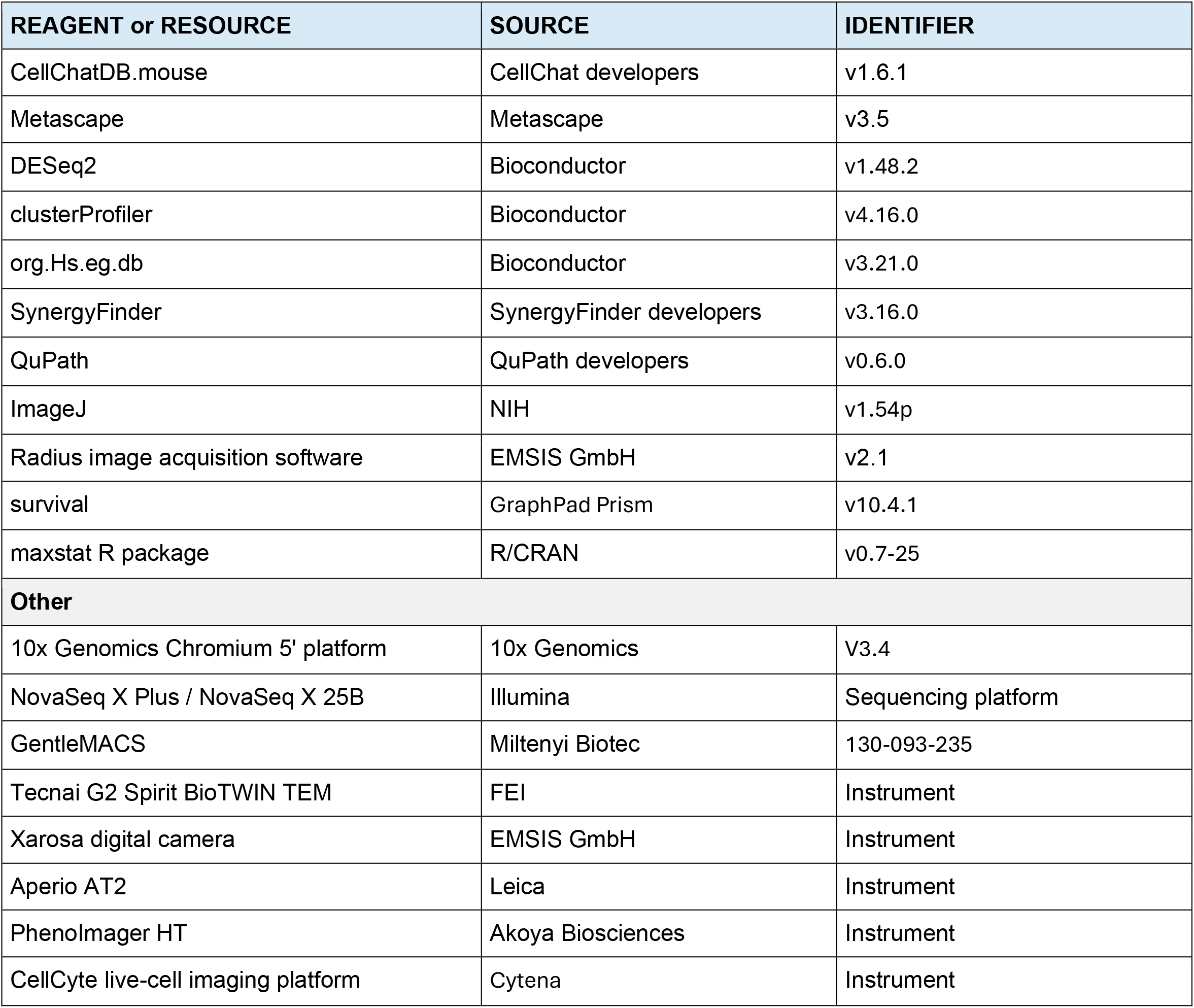

### RESOURCE AVAILABILITY

#### Lead contact

Further information and requests for resources and reagents should be directed to and will be fulfilled by the lead contact

#### Materials availability

This study did not report the generation of new unique reagents in the supplied Methods. Patient-derived glioblastoma stem cell (GSC) models were obtained through the Mayo Clinic BRIDGE Biobank under the approvals described below. Availability of these models and any other study materials is subject to institutional approvals and applicable material-transfer requirements.

#### Data and code availability

Bulk RNA-sequencing, single-cell RNA-sequencing, and associated processed datasets are described in this study. Any additional information required to reanalyze the data reported in this paper will be made available by the lead contact upon reasonable request, subject to applicable institutional and data-use restrictions.

### EXPERIMENTAL MODEL AND STUDY PARTICIPANT DETAILS

#### Human glioblastoma specimens and patient-derived GSC models

Human glioblastoma (GBM) specimens were obtained with informed consent through the Mayo Clinic BRIDGE Biobank under Institutional Review Board-approved protocols 16-008485 and 17-003013. Fresh tumor specimens were dissociated into single-cell suspensions and established as patient-derived GSC cultures as previously described. The models used in this study were GBM1A, QNS690, QNS712, and QNS986. GSCs were maintained in serum-free DMEM/F12 supplemented with EGF, FGF, and antibiotics on laminin-coated culture vessels at 37°C in 5% CO2. Experiments were performed using cultures below passage 15.

#### Orthotopic patient-derived GBM xenograft models

All animal studies were approved by the institutional IACUC under protocol A00008191-25. GFP/luciferase-expressing GBM1A or QNS690 GSCs (5 x 10^5 cells) were implanted intracranially into Nude mice to establish orthotopic GBM xenografts. Tumor growth was monitored by bioluminescence imaging. Following tumor establishment, animals received vehicle or clemastine plus bexarotene using the systemic or intracranial regimens described below.

#### Syngeneic CT-2A GBM model

To evaluate treatment effects in an immunocompetent setting, syngeneic orthotopic tumors were established by intracranial implantation of 5 x 10^4 CT-2A GFP-Luc cells into C57BL/6 mice. After tumor engraftment was confirmed, animals received an intracranial Ommaya-like reservoir and were treated with vehicle or clemastine plus bexarotene. Animals used for early mechanistic and immune analyses were euthanized 72 h after treatment initiation.

## METHOD DETAILS

### Patient-derived GSC culture and drug treatment

GBM1A, QNS690, QNS712, and QNS986 GSCs were cultured in serum-free DMEM/F12 containing EGF, FGF, and antibiotics on laminin-coated vessels at 37°C and 5% CO2. Clemastine (Clem) and bexarotene (Bex) were prepared in DMSO. Unless otherwise specified, cells were treated for 72 h with vehicle (DMSO), Clem (12 μM), Bex (30 μM), or the combination. Dose-response and immunoblot experiments used the concentrations indicated below.

### Cell viability, proliferation, and drug-synergy assays

Drug sensitivity was determined by exposing patient-derived GSCs to increasing concentrations of Clem or Bex (0.1-50 μM) for 72 h. Cell abundance was measured using the CyQUANT Cell Proliferation Assay according to the manufacturer’s instructions, and half-maximal inhibitory concentrations (IC50) were estimated by nonlinear regression. Each condition was tested in quadruplicate and experiments were repeated independently. For longitudinal growth analysis, GSCs were treated with vehicle, Clem (3-12 μM), Bex (10-30 μM), or the corresponding dose combinations and imaged every 4 h for 7 days using a CellCyte live-cell imaging platform. Growth was quantified from confluence measurements and normalized to baseline. Drug interaction at 72 h was evaluated in SynergyFinder using normalized viability data. Highest Single Agent (HSA) synergy scores were used for the synergy analyses represented in the supplementary figures.

### Western blotting

Patient-derived GSCs were treated for 72 h with vehicle (DMSO), Clem (6 or 12 μM), Bex (20 or 30 μM), or the corresponding combinations. Cells were lysed in RIPA buffer supplemented with protease and phosphatase inhibitors, and protein concentration was determined by BCA assay. Equal amounts of protein were separated by SDS-PAGE and transferred to PVDF membranes. Membranes were incubated with primary antibodies against stemness, cholesterol-metabolism, ER-stress/UPR, and autophagy proteins, followed by HRP-conjugated secondary antibodies. Signal was detected by enhanced chemiluminescence. Band intensities were quantified in ImageJ and normalized to the corresponding loading control. Experiments were independently repeated.

### Immunocytochemistry and image quantification

Patient-derived GSCs were seeded in laminin-coated 96-well plates at 5 x 10^3 cells per well and allowed to attach overnight. Cells were treated with vehicle (DMSO), Clem (12 μM), Bex (30 μM), or Clem plus Bex for 72 h, fixed with 4% paraformaldehyde for 20 min at room temperature, permeabilized with Triton X-100 (0.1% for cytoplasmic targets or 0.25% for nuclear targets), and blocked with 5% normal goat serum for 1 h. Primary antibodies were incubated overnight at 4°C in blocking buffer: Ki67 (mouse, 1:100), cleaved caspase-3 (mouse, 1:100), Nestin (mouse, 1:200), GFAP (rabbit, 1:500), SOX2 (rat, 1:200), Vimentin (rabbit, 1:100), DHCR7 (rabbit, 1:100), EBP (mouse, 1:100), FABP7 (rabbit, 1:100), APOE (mouse, 1:200), LDLR (rabbit, 1:100), ABCA1 (mouse, 1:50), phospho-IRE1 S724 (rabbit, 1:100), SQSTM1/p62 (mouse, 1:150), LAMP1 (mouse, 1:200), and IDOL (rabbit, 1:20). Cells were then incubated with the appropriate fluorophore-conjugated secondary antibodies for 1 h at room temperature and counterstained with DAPI. Confocal images were acquired from three random 10x fields per well, with three wells per condition (nine images per condition). Protein expression and cell number were quantified using QuPath, with cell counts based on DAPI-positive nuclei. Experiments were performed in triplicate and repeated independently.

### Bulk RNA sequencing and pathway analysis

Bulk RNA-sequencing data from patient-derived GSC models treated with vehicle, Clem, Bex, or the combination were analyzed with DESeq2. Differential-expression results and normalized expression matrices were exported for downstream analysis. For pathway-focused heatmaps, normalized expression values were log2 transformed and scaled by gene using z-score transformation. Gene set enrichment analysis (GSEA) used preranked gene lists derived from DESeq2 Wald statistics. Gene Ontology Biological Process enrichment was performed with clusterProfiler and the org.Hs.eg.db annotation database using gseGO, with minimum and maximum gene-set sizes of 10 and 500, respectively, and Benjamini-Hochberg correction for multiple testing. Enrichment plots displayed normalized enrichment score (NES), gene-set size, and adjusted P value, with common bubble-size and color scales maintained for comparative visualization.

### Transmission electron microscopy

Patient-derived GSCs were seeded in 8-well Permanox chamber slides and treated with vehicle (DMSO), Clem (12 μM), Bex (30 μM), or Clem plus Bex for 72 h. Cells were fixed in 3.5% glutaraldehyde and 1% paraformaldehyde in PBS, post-fixed in 2% osmium tetroxide, dehydrated through graded ethanol, and embedded in Durcupan resin. Ultrathin 70-nm sections were cut with a diamond knife and examined on a Tecnai G2 Spirit BioTWIN transmission electron microscope equipped with a Xarosa digital camera. Images were acquired using Radius v2.1. Ultrastructural features associated with intracellular lipid accumulation, lipid droplets, endoplasmic-reticulum stress, and autophagic vesicles were evaluated across treatment conditions.

### BODIPY-cholesterol uptake and accumulation assay

Patient-derived GSCs were seeded on laminin-coated chamber slides or 96-well plates and allowed to attach overnight. Cells were incubated with fluorescent BODIPY-cholesterol according to the manufacturer’s recommendations for 2 h to permit incorporation into cellular cholesterol pools. Cells were then treated with vehicle (DMSO), Clem (12 μM), Bex (30 μM), or the combination for 72 h. Images were acquired every 2 h for 72 h to monitor intracellular cholesterol accumulation and cholesterol-rich lipid-droplet formation.

### Orthotopic GBM implantation and tumor monitoring

GFP/luciferase-expressing GBM1A or QNS690 GSCs (5 x 10^5 cells) were implanted intracranially into Nude mice. Tumor engraftment and progression were followed by bioluminescence imaging. Following establishment of detectable tumors, mice were assigned to vehicle or Clem plus Bex treatment. For mechanistic analyses, brains were collected 72 h after treatment for histology, immunofluorescence, and single-cell RNA sequencing. Survival cohorts were followed for tumor progression and analyzed by Kaplan-Meier methods.

### Systemic and intracranial Clem+Bex administration

For systemic treatment, tumor-bearing mice received Clem at 7.5 mg/kg plus Bex at 30 mg/kg by intraperitoneal injection 5 days per week for 6 weeks. For local treatment, an Ommaya-like intracranial reservoir was implanted and vehicle or Clem plus Bex was administered every 3 days for 6 weeks at the intracranial doses specified for the corresponding experiment. The supplied Methods did not provide a single numeric intracranial concentration applicable to all experiments; therefore, dose values should be reported with the relevant experiment or figure.

### Syngeneic CT-2A model and peripheral immune profiling

C57BL/6 mice bearing intracranial CT-2A GFP-Luc tumors received an Ommaya-like intracranial reservoir after confirmation of tumor engraftment and were treated with vehicle or Clem plus Bex using the intracranial regimen employed for the patient-derived orthotopic studies. For early mechanistic studies, animals were euthanized 72 h after treatment initiation. Brains and peripheral tissues were collected for histology and single-cell transcriptomic analyses. Peripheral leukocyte populations were evaluated from triplicate blood smears for each animal (n=6 mice per condition) and quantified manually.

### Single-cell library preparation and sequencing

Tumors were mechanically and enzymatically dissociated into single-cell suspensions using GentleMACS according to the manufacturer’s protocol. Single-cell libraries were generated using the 10x Genomics Chromium 5′ platform according to the manufacturer’s instructions. Gene-expression libraries were sequenced on an Illumina NovaSeq X Plus using NovaSeq X 25B reagents to a target depth of approximately 50,000 reads per cell.

### Mouse tissue collection, histology, and immunohistochemistry

At experimental endpoints, mice were perfused with PBS followed by 4% paraformaldehyde. Brains were post-fixed overnight, paraffin embedded, sectioned, and processed for histology. H&E staining was used to assess tumor morphology. For immunohistochemistry, sections underwent pH 6 antigen retrieval and were stained with mouse anti-HuNu (1:200) or rabbit anti-GFP (1:300), followed by DAB detection. Whole-slide images were acquired using an Aperio AT2 scanner, and tumor cells were quantified using eSlide Manager.

### Akoya multiplex immunofluorescence

Multiplex immunofluorescence was performed using the Akoya Opal workflow. Orthotopic xenograft tumors were stained for SQSTM1/p62 (mouse, 1:400; AR6), HuNu (mouse, 1:200; AR6), and BiP (rabbit, 1:100; AR6). Syngeneic tumors were stained for Iba1 (rabbit, 1:4000; AR9), Arg1 (rabbit, 1:100; AR6), CD4 (rabbit, 1:50; AR6), CD8 (rabbit, 1:200; AR6), and CD19 (rabbit, 1:800; AR6). Opal tyramide signal amplification was used with antibody stripping between staining cycles. Slides were counterstained with DAPI and mounted. Multispectral images were acquired using an Akoya PhenoImager HT under matched exposure settings across treatment groups. Spectral unmixing, tissue segmentation, and cell phenotyping were performed in QuPath. Depending on the analysis, marker abundance was reported as percentage of positive cells, marker-positive area, or fluorescence intensity within defined tumor regions.

### Single-cell RNA-sequencing computational analysis

Single-cell RNA-sequencing data were processed with Cell Ranger v9.0.1. For mouse brains bearing human tumors, reads were aligned to a custom combined human GRCh38/mouse GRCm39 reference. Ambient RNA contamination was removed with decontX from the celda package, and cells were classified as human or mouse when more than 50% of transcripts were species specific. Quality-control thresholds were optimized separately for human and mouse cells. Downstream analyses were performed in Seurat v5.4.0. Batch effects were corrected with Harmony, clustering and visualization were performed using UMAP, and cell types were annotated from established marker genes. Differential expression was assessed with MAST and pathway enrichment with Metascape v3.5. For syngeneic tumors, reads were aligned to GRCm39, ambient RNA was removed with decontX, and cells passing quality control were analyzed with the same Seurat, Harmony, MAST, and Metascape workflow.

### CellChat analysis

Cell-cell communication was inferred from single-cell RNA-sequencing data using CellChat v1.6.1 and the CellChatDB.mouse ligand-receptor database. Communication probabilities and signaling pathways were calculated using default parameters, and interactions supported by fewer than 10 cells were excluded. Global networks were compared between treatment groups using interaction counts, information flow, and rankNet analyses. Pathway-specific analyses focused on MIF, MHC-I, NKG2D, CXCL, and CD96 signaling to identify dominant ligand-receptor interactions and the relative contributions of cell populations as senders, receivers, mediators, and influencers.

### GBM cellular-state and cell-cycle analysis

Single-cell RNA-sequencing data from GBM1A, QNS690, QNS712, and QNS986 GSC models treated with vehicle or Clem (12 μM) plus Bex (30 μM) were analyzed in R. Filtered Cell Ranger matrices were scored using published Neftel transcriptional-state signatures representing AC-like, MES-like, NPC-like, and OPC-like programs, together with G1/S and G2/M cycling signatures. Cells were classified as cycling according to their G1/S and G2/M scores and visualized using Neftel butterfly plots. The proportions of cycling and non-cycling cells within each transcriptional state were quantified across treatment groups.

### TCGA pathway-signature and survival analysis

Normalized TCGA GBM RNA-sequencing expression data (FPKM) and survival information were analyzed to evaluate cholesterol, ER/UPR, and autophagy-related signatures. Genes enriched in the relevant pathways were identified from GSEA, and pathway scores were generated from the first principal component (PC1) of scaled gene expression. Patients were stratified into high- and low-signature groups using maximally selected rank statistics. Overall survival was evaluated by Kaplan-Meier analysis, log-rank testing, and univariate Cox proportional-hazards regression using the survival and maxstat R packages. Statistical significance was defined as P < 0.05.

## QUANTIFICATION AND STATISTICAL ANALYSIS

Quantitative analyses were performed using the statistical approaches specified for each experiment. Immunocytochemistry experiments used three random 10x fields from each of three wells per condition (nine images per condition), with experiments performed in triplicate and repeated independently. Peripheral blood-smear analyses in the CT-2A study included six animals per condition and triplicate smears per animal. For immunofluorescence quantification shown in the supplementary analyses, data were reported as mean ± SEM and group comparisons were performed using one-way ANOVA with multiple-comparisons correction (Tukey’s test where specified). Drug-synergy analyses used normalized 72-h viability values and HSA scoring. Bulk RNA-seq differential expression was analyzed with DESeq2, and pathway enrichment used Benjamini-Hochberg-adjusted P values. Single-cell differential expression was assessed with MAST. Survival analyses used Kaplan-Meier curves, log-rank tests, and univariate Cox regression. Unless otherwise stated, P < 0.05 was considered statistically significant.

Significance notation used in the associated figures was *P < 0.05, **P < 0.01, ***P < 0.001, and ****P < 0.0001 where applicable. Exact n values, the definition of n (biological versus technical replicate), exact P values, and the statistical test used for each figure should be reported in the corresponding figure legends or source-data files.

## Acknowledgements

We thank Marina Hanson, DVM, PhD, and the Department of Comparative Medicine at Mayo Clinic Florida for assistance with the intracranial Ommaya reservoir model; Brandy Edenfield for histology support; and the Mayo Clinic BRIDGE Biobank for providing clinically annotated GBM specimens. We also thank Mario Soriano-Navarro and the Microscopy Core at the Centro de Investigación Príncipe Felipe for electron microscopy support. Finally, we are deeply grateful to the patients and their families for generously donating tissue for this research

## Funding

National Cancer Institute, National Institutes of Health grant K99CA318464-01 (M.J.U.-N.).

Center for Innovation in Brain Tumor Therapeutics (M.J.U.-N., A.Q.-H.).

Mayo Clinic Clinician Investigator Award (A.Q.-H.).

William J. and Charles H. Mayo Named Professorship (A.Q.-H.).

Monica Flynn Jacoby Endowed Chair (A.Q.-H.). Uihlein Neuro-Oncology Research Fund (A.Q.-H.).

Florida Department of Health Cancer Research Chair Fund (A.Q.-H.).

Jacquie Lorraine Goldman Fund for Brain Tissue Bank (A.Q.-H.).

Valencian Council for Education, Culture, University, and Employment grant CIPROM/2023/053 (V.H.-P.)

## .Author Contributions

M.J.U.-N. conceived and designed the study, performed experiments, analyzed and interpreted data, supervised research activities, generated figures, and wrote the manuscript. R.M.W., V.K.J., M.L., M.B., A.N.B., C.G.-P., E.S.-G., P.S., J.P.N.G.L., and B.T. performed experiments and contributed to data acquisition and analysis. A.N., A.R.-M., and Y.R. performed bioinformatic analyses and contributed to data interpretation. R.M.-G., V.H.-P., and J.M.G.-V. performed and interpreted electron microscopy studies. S.R., V.E.C., L.D., H.D., and H.Q. provided scientific supervision, technical expertise, and interpretation of results. A.Q.-H. conceived and supervised the study, secured funding, interpreted data, and edited the manuscript. All authors reviewed, edited, and approved the final manuscript.

