## Supplementary material for "Intracranial Targeting of Cholesterol Processing Reveals a Therapeutic Vulnerability that Reprograms Glioblastoma and Promotes Antitumor Immunity": Supp figs

### SUPPLEMENTARY FIGURES

#### Supplementary Figure 1

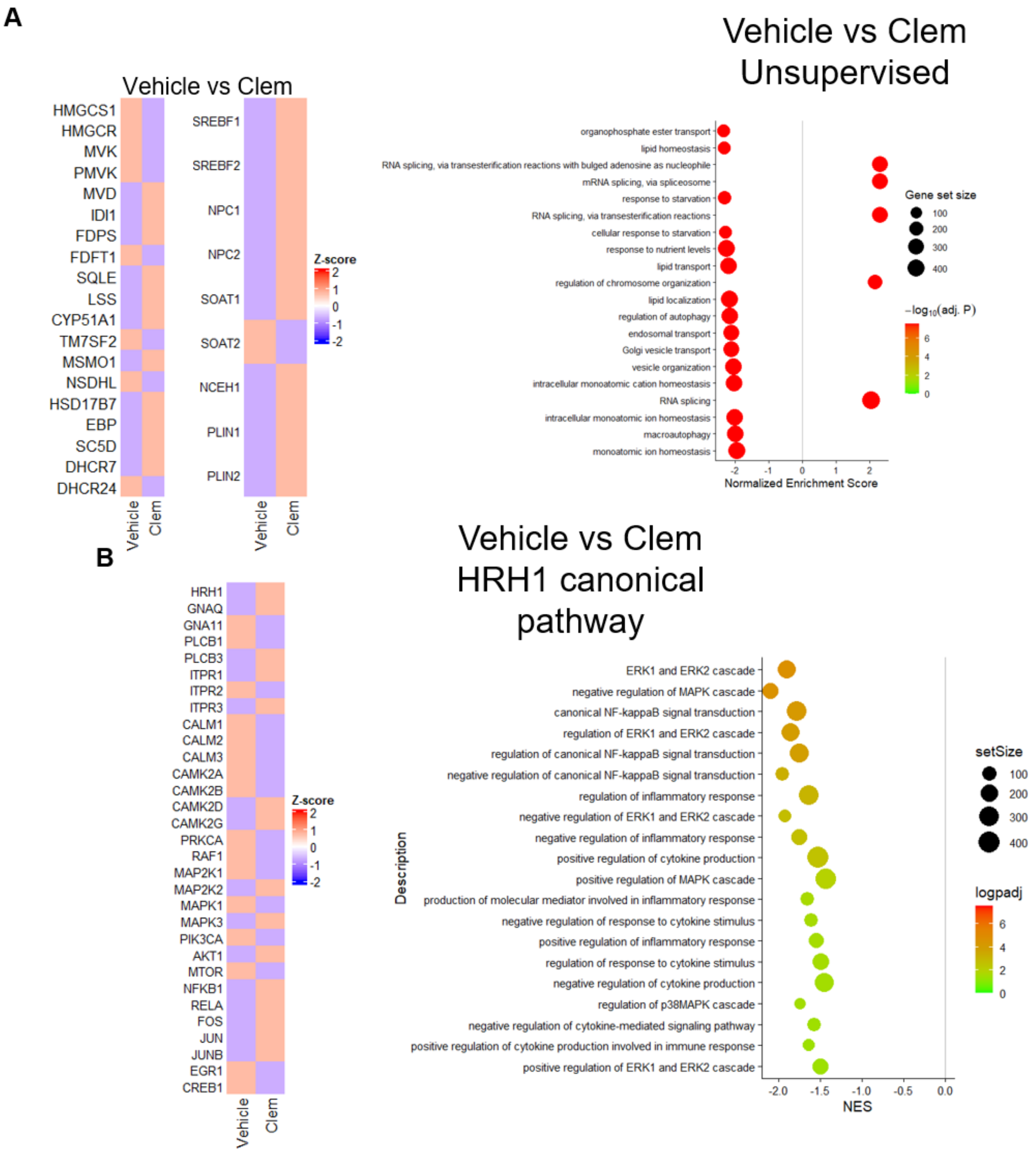


**Supplementary Figure 1. Clem primarily alters cholesterol metabolism rather than canonical HRH1 signaling in GSCs. A)** Left, heatmap of cholesterol biosynthesis (HMGCS1–DHCR24) and cholesterol storage/trafficking (SREBF1–PLIN2) genes in vehicle- and clemastine-treated glioblastoma stem cells (GSCs), averaged across four patient-derived lines. Right, unsupervised GO Biological Process GSEA showing enrichment of pathways related to lipid homeostasis, transport, vesicular trafficking, and autophagy. Positive NES indicates enrichment in clemastine-treated cells. **B)** Left, heatmap of canonical histamine H1 receptor (HRH1) signaling genes in vehicle- and clemastine-treated GSCs. Right, enrichment analysis of curated HRH1-related pathways. Compared with the extensive alterations observed in cholesterol metabolism, canonical HRH1 signaling exhibited only modest transcriptional changes, supporting cholesterol dysregulation as the predominant response to clemastine treatment.

Heatmaps show row-wise Z-score–scaled expression from bulk RNA-seq data. Bubble size indicates gene set size and color indicates –log10(adjusted P value).

#### Supplementary Figure 2

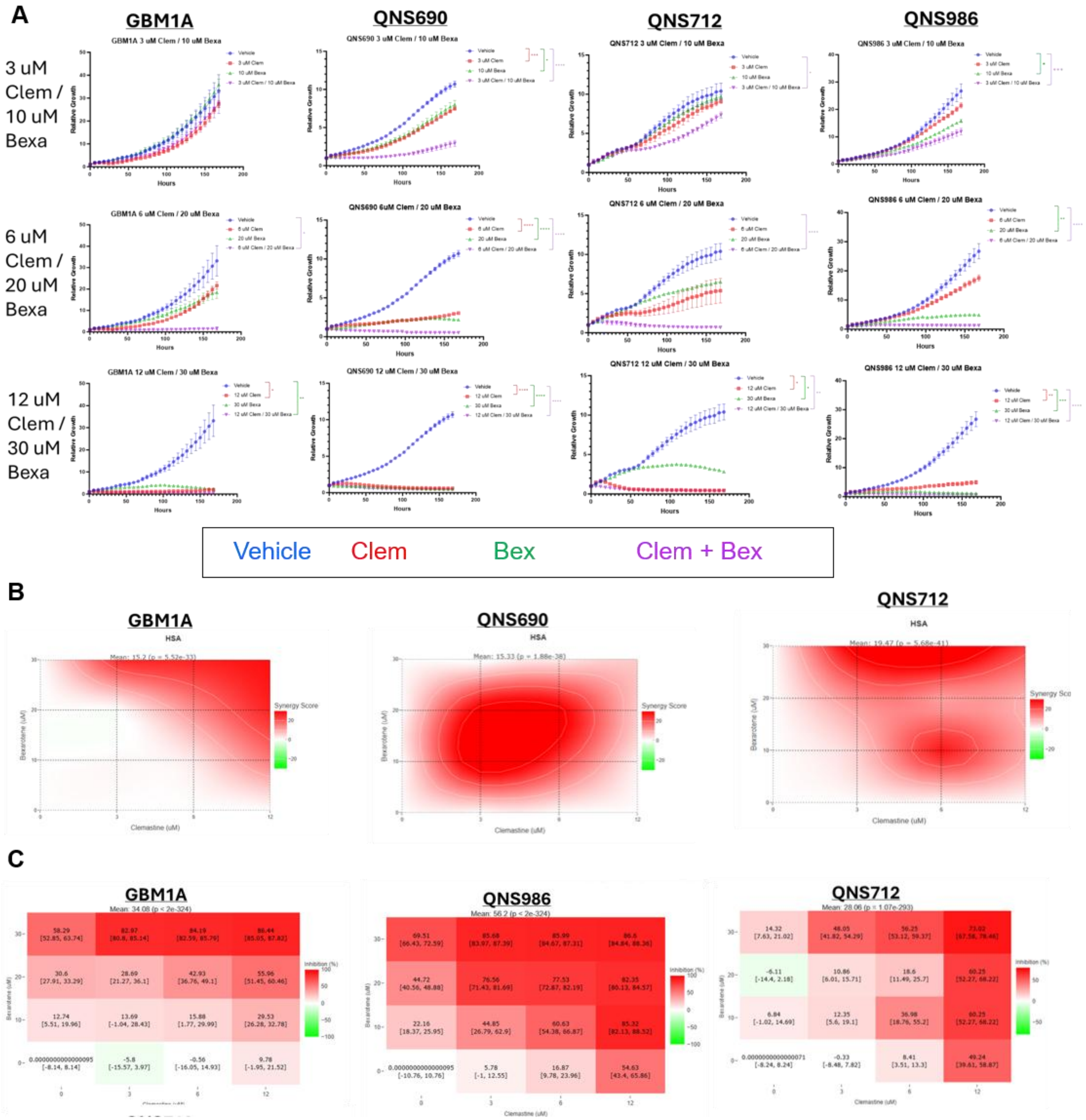


**Supplementary Figure 2. Clemastine+Bexarotene treatment synergistically impairs GSC expansion and in patient-derived GBM models. A)** Growth curves of four primary GBM cell lines treated with vehicle, Clem Bex, or the combination. Combination treatment at IC~50 produced the strongest inhibition of cell expansion and significantly reduced proliferation compared with either monotherapy, demonstrating synergistic antitumor activity across molecularly distinct GBM models. *P < 0.05, **P < 0.01, ***P < 0.001, ****P < 0.0001. **B)** Representative HSA synergy maps and **C)** growth inhibition matrices across all patient-derived GSC lines treated with increasing concentrations of Clem and Bex. Positive HSA scores indicate synergistic interactions, with robust synergy observed across multiple dose combinations in all models. Growth inhibition analyses demonstrated significantly greater suppression of tumor cell viability with combination treatment compared with either agent alone, confirming cooperative antitumor effects across genetically distinct GBM cultures. Values represent mean response with 95% confidence intervals. HSA, Highest Single Agent.

**Supplementary Figure 3**

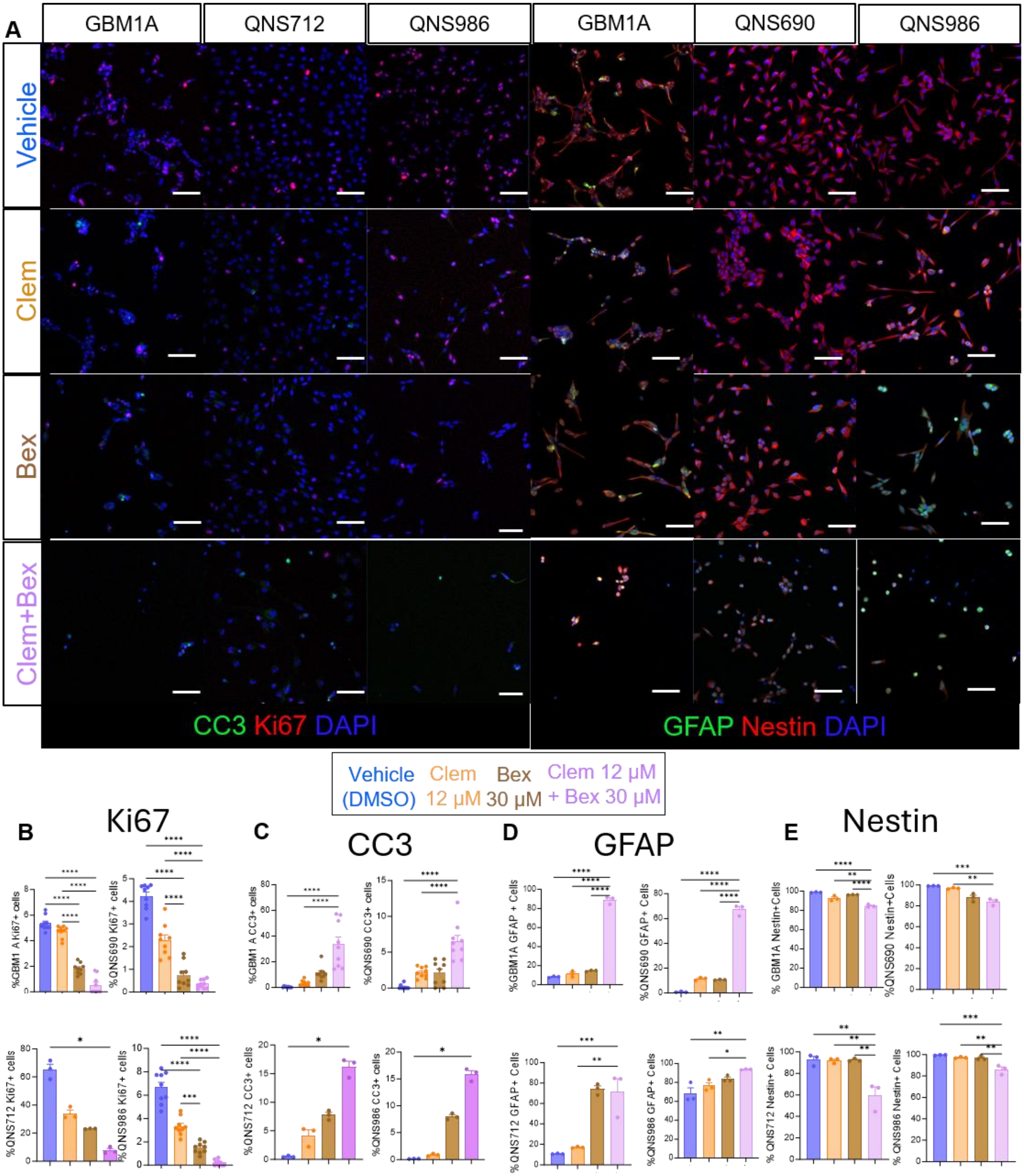


**Supplementary Figure 3. Clemastine+Bexarotene promotes apoptosis and differentiation while reducing stemness in GBM GSCs. A)** Representative immunofluorescence images of CC3, Ki67, GFAP, and Nestin in GBM1A, QNS690, QNS712, and QNS986 GSCs treated with vehicle, Clem (12 µM), Bex (30 µM), or the combination. Scale bars, 100 µm. **B–E)** Quantification of Ki67-positive (B), CC3-positive (C), GFAP-positive (D), and Nestin-positive (E) cells. Data are mean ± SEM. One-way ANOVA with Tukey's multiple-comparisons test. *P < 0.05, **P < 0.01, ***P < 0.001, ****P < 0.0001

**Supplementary Figure 4**

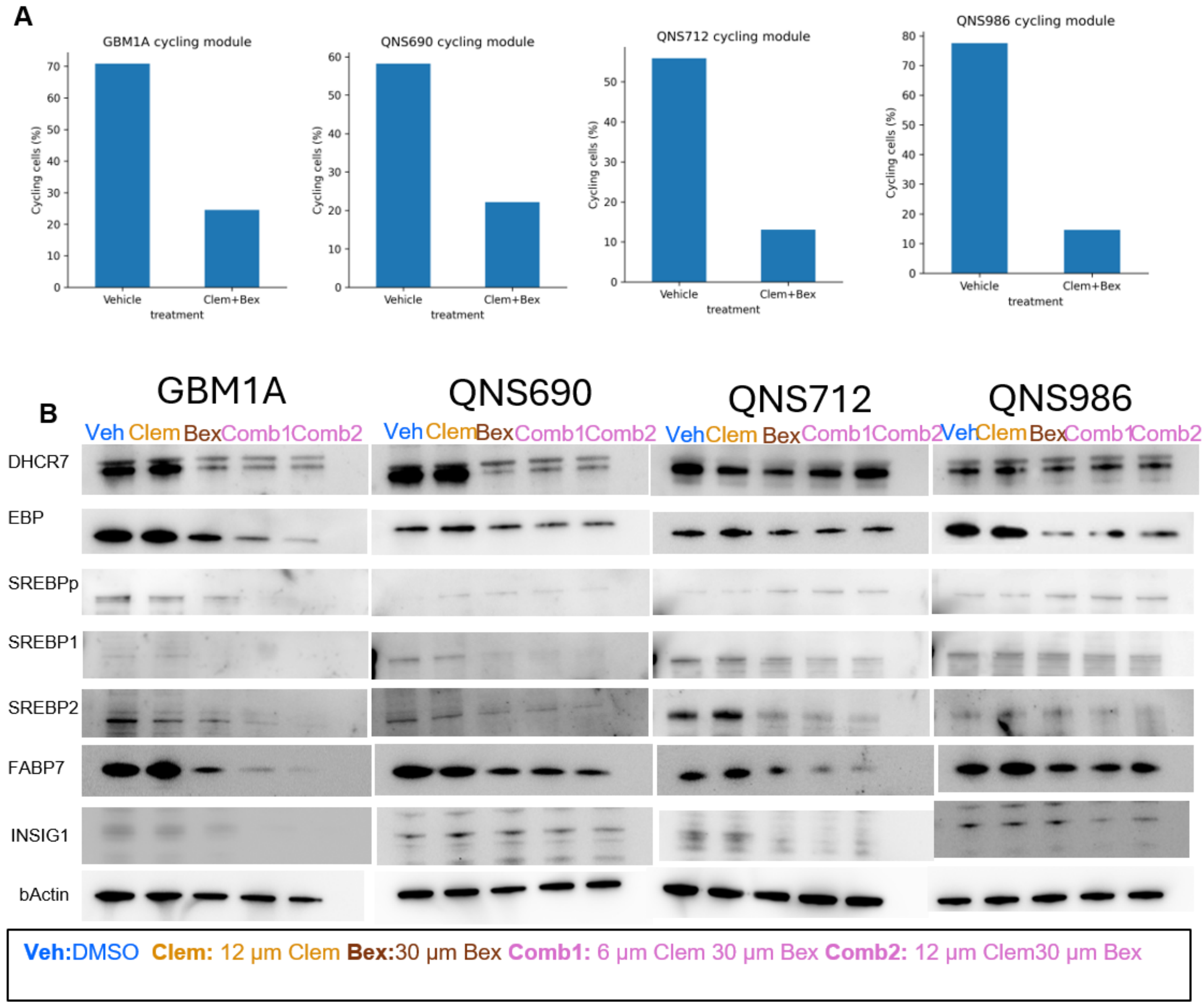


**Supplementary Figure 4. Clemastine and bexarotene suppress proliferative programs and alter cholesterol homeostasis regulators in GSCs. A)** Quantification of cycling cells in four patient-derived GSC models (GBM1A, QNS690, QNS712, and QNS986) following treatment with vehicle or Clem+Bex. Cycling cells were identified using G1/S and G2/M cell-cycle gene modules derived from single-cell RNA sequencing datasets. Across all four models, combination treatment markedly reduced the proportion of cycling cells, consistent with suppression of tumor proliferation. **B)** Representative western blot analysis of cholesterol biosynthesis and regulatory proteins in GBM1A, QNS690, QNS712, and QNS986 cells treated with vehicle (DMSO), Clem (12 µM), Bex (30 µM), or two combination regimens (Comb1: 6 µM clemastine + 20 µM bexarotene; Comb2: 12 µM clemastine + 30 µM bexarotene). Expression of DHCR7, EBP, precursor SREBP (SREBPp), SREBP1, SREBP2, FABP7, and INSIG1 was assessed by immunoblotting. Combination treatment resulted in broad alterations of cholesterol biosynthesis and lipid regulatory pathways, including reduced expression of cholesterol synthesis enzymes and modulation of SREBP signaling. β-actin served as a loading control.

#### Supplementary Figure 5

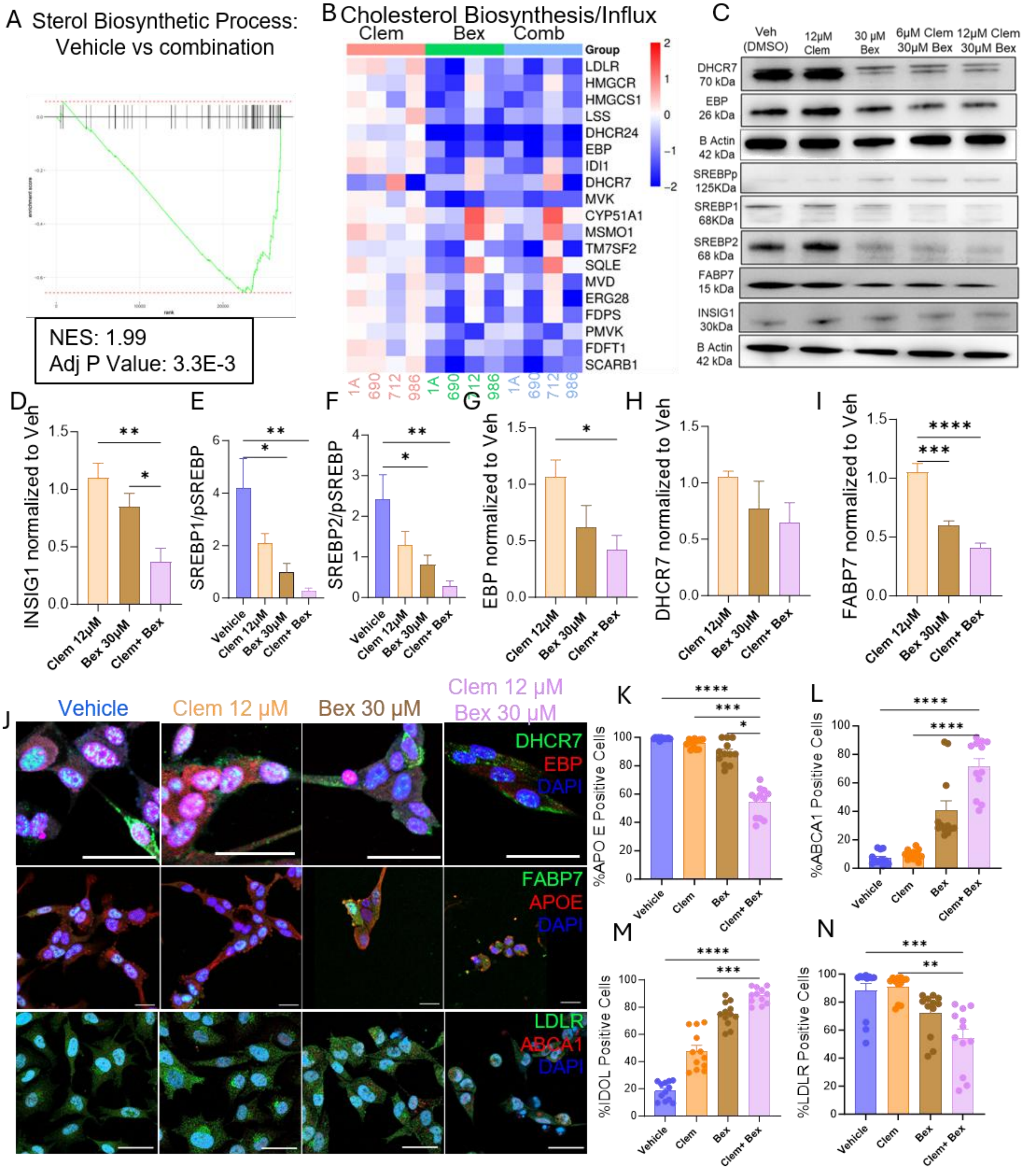

**Supplementary Figure 5. Clemastine+Bexarotene disrupts cholesterol biosynthesis and transport in GBM. A)** GSEA showing enrichment of sterol biosynthesis pathways following combination treatment. **B)** Heatmap of cholesterol biosynthesis and transport genes across patient-derived GBM models. **C)** Western blots of DHCR7, EBP, SREBP1, SREBP2, FABP7, and INSIG1. **D–I)** Quantification of INSIG1, SREBP1, SREBP2, EBP, DHCR7, and FABP7. **J)** Representative immunofluorescence images of DHCR7, EBP, FABP7, APOE, LDLR, and ABCA1. Scale bars, 100 µm (DHCR7, EBP) and 25 µm (FABP7, APOE, LDLR, ABCA1). **K–N)** Quantification of APOE-, ABCA1-, IDOL-, and LDLR-positive cells, demonstrating reduced cholesterol influx and increased efflux following Clem+Bex treatment. Data are mean ± SEM. One-way ANOVA with Tukey's multiple-comparisons test. *P < 0.05, **P < 0.01, ***P < 0.001, ****P < 0.0001.

**Supplementary Figure 6**

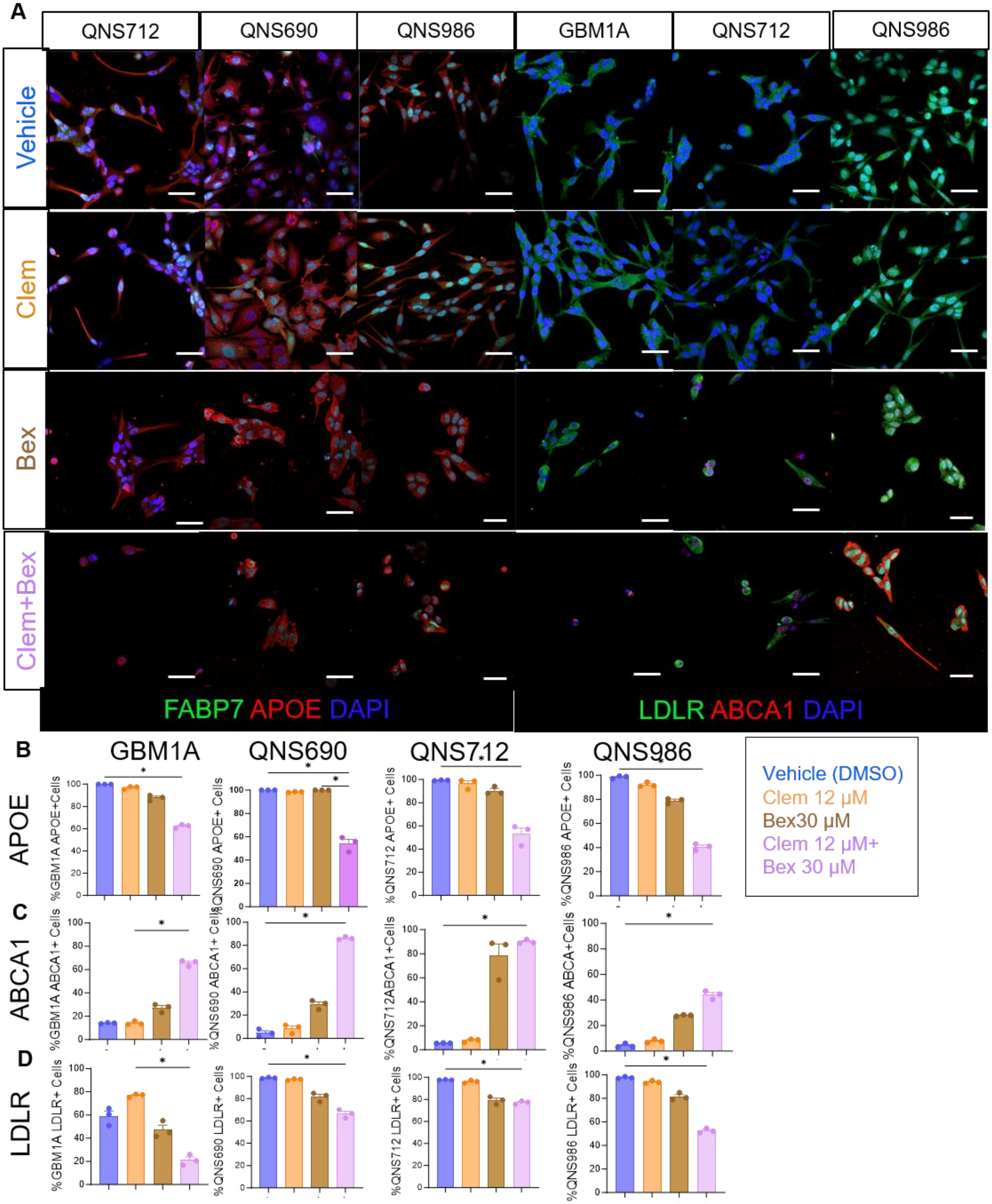


**Supplementary Figure 6. Clemastine+Bexarotene remodels cholesterol transport in patient-derived GSCs. A)** Representative immunofluorescence images of FABP7, APOE, LDLR, and ABCA1 following treatment. Scale bars, 25 μm. **B–D)** Quantification of APOE-, ABCA1-, and LDLR-positive cells, demonstrating reduced cholesterol uptake and increased efflux after combination therapy. Data are mean ± SEM. One-way ANOVA with multiple-comparisons correction. *$P < 0.05$.

#### Supplementary Figure 7

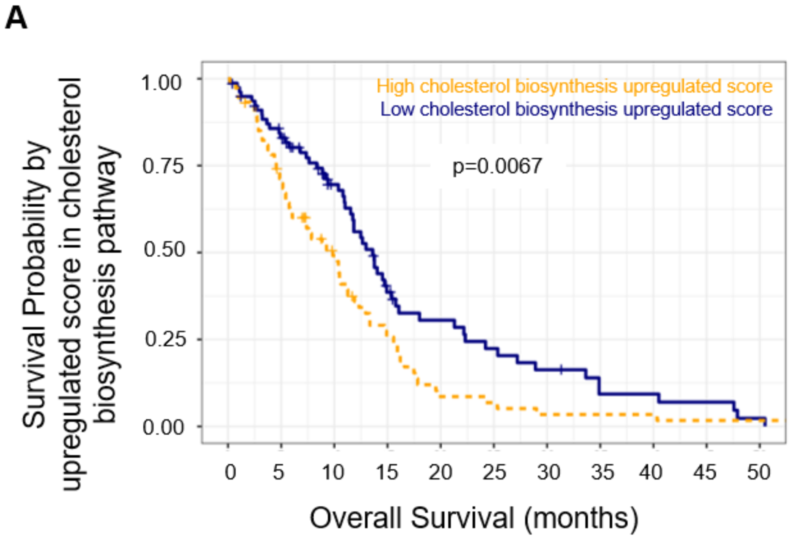


**Supplementary Figure 7. Cholesterol biosynthesis pathway activation correlates with poor survival in GBM. A)** Kaplan–Meier survival analysis of TCGA GBM patients stratified by cholesterol biosynthesis pathway activation score. Patients with elevated cholesterol biosynthesis pathway activation demonstrated significantly increased overall survival compared with patients with low pathway activation.

#### Supplementary Figure 8

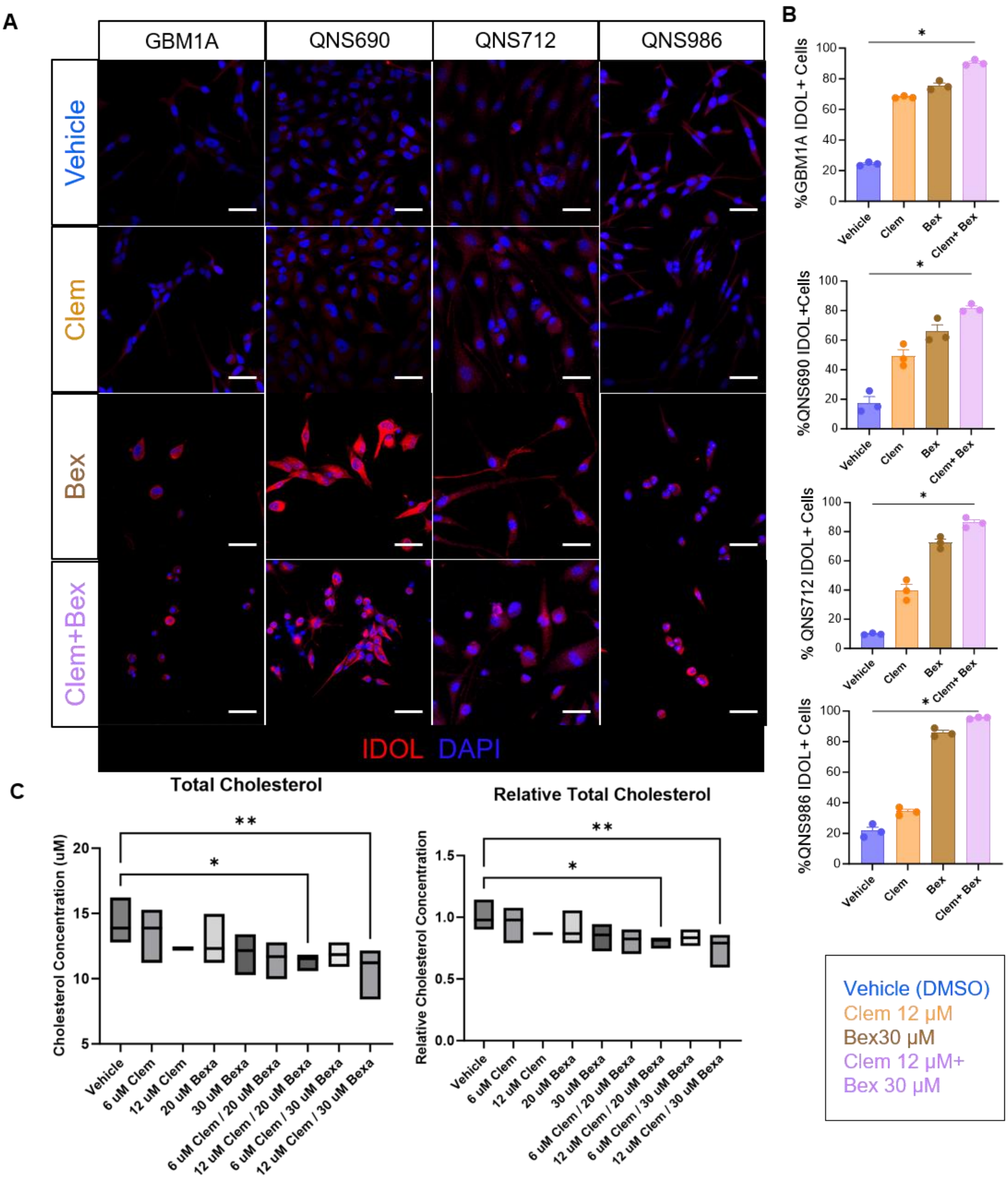


**Supplementary Figure 8. Clemastine+Bexarotene increases IDOL expression and reduces intracellular cholesterol. A)** Representative IDOL immunofluorescence images of GBM1A, QNS690, QNS712, and QNS986 following treatment. Scale bars, 50 µm. **B)** Quantification of IDOL-positive cells. **C)** Bioluminescent quantification of intracellular cholesterol. Data are mean ± SEM. *P < 0.05.

**Supplementary Figure 9**

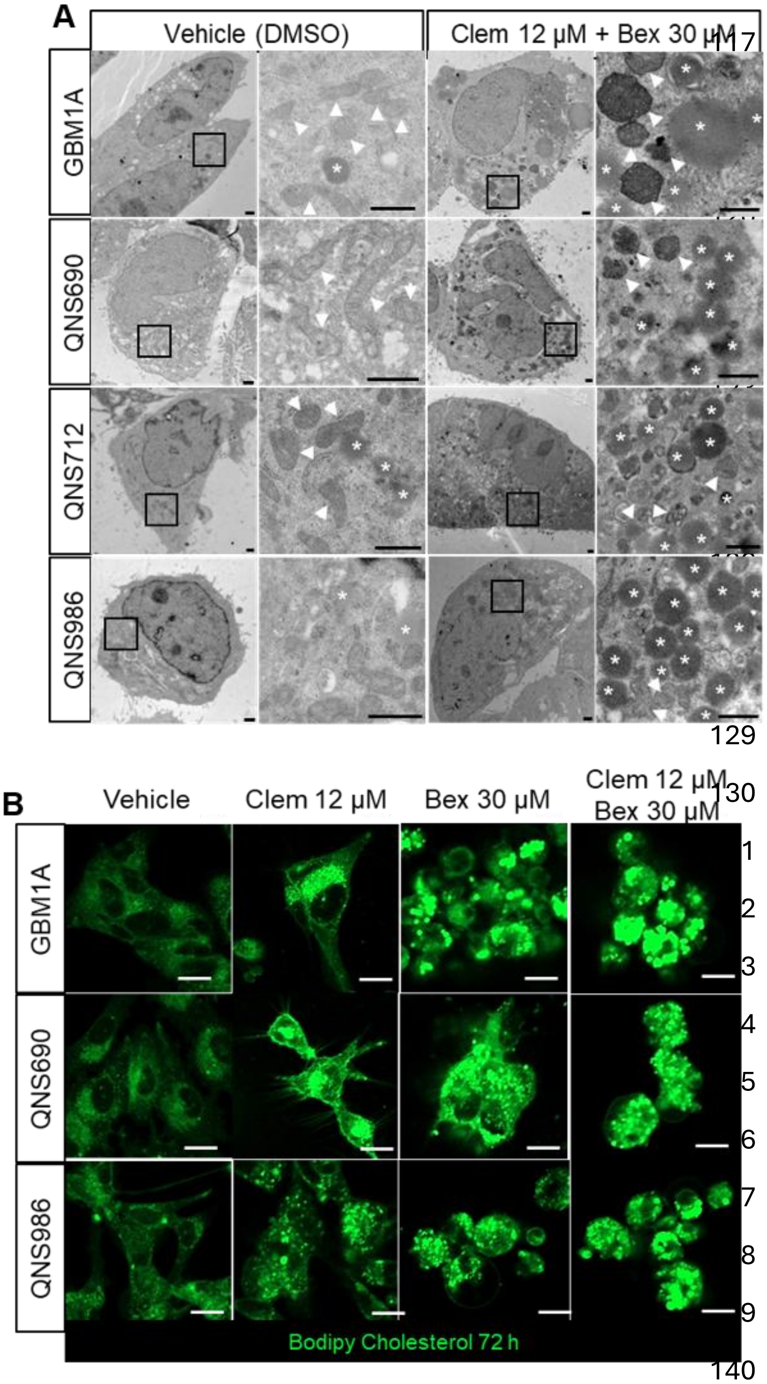


**Supplementary Figure 9. Clemastine and bexarotene induce intracellular cholesterol accumulation and lipid droplet formation in patient-derived GSCs. A)** Transmission electron microscopy (TEM) of GBM1A, QNS690, QNS712, and QNS986 cells treated with vehicle (DMSO) or Clem + Bex for 72 h. Vehicle-treated cells displayed normal cellular ultrastructure with sparse lipid storage. In contrast, combination-treated cells accumulated abundant electron-dense lipid droplets (*) and exhibited extensive intracellular lipid deposition, consistent with disrupted cholesterol trafficking and storage. Arrowheads indicate mitochondria. Scale Bars: 1 μm. **B)** Representative BODIPY-cholesterol staining of GBM1A, QNS690, and QNS986 cells following treatment with vehicle, Clem, Bex, or the combination for 72 h. Treatment increased intracellular cholesterol accumulation, with the strongest effect observed in combination-treated cells, resulting in prominent cholesterol-rich lipid droplets. Together, these findings demonstrate that clemastine and bexarotene induce profound alterations in cholesterol homeostasis, leading to intracellular cholesterol sequestration and lipid droplet accumulation in GBM cells. Scale Bars: 50 μm.

#### Supplementary Figure 10

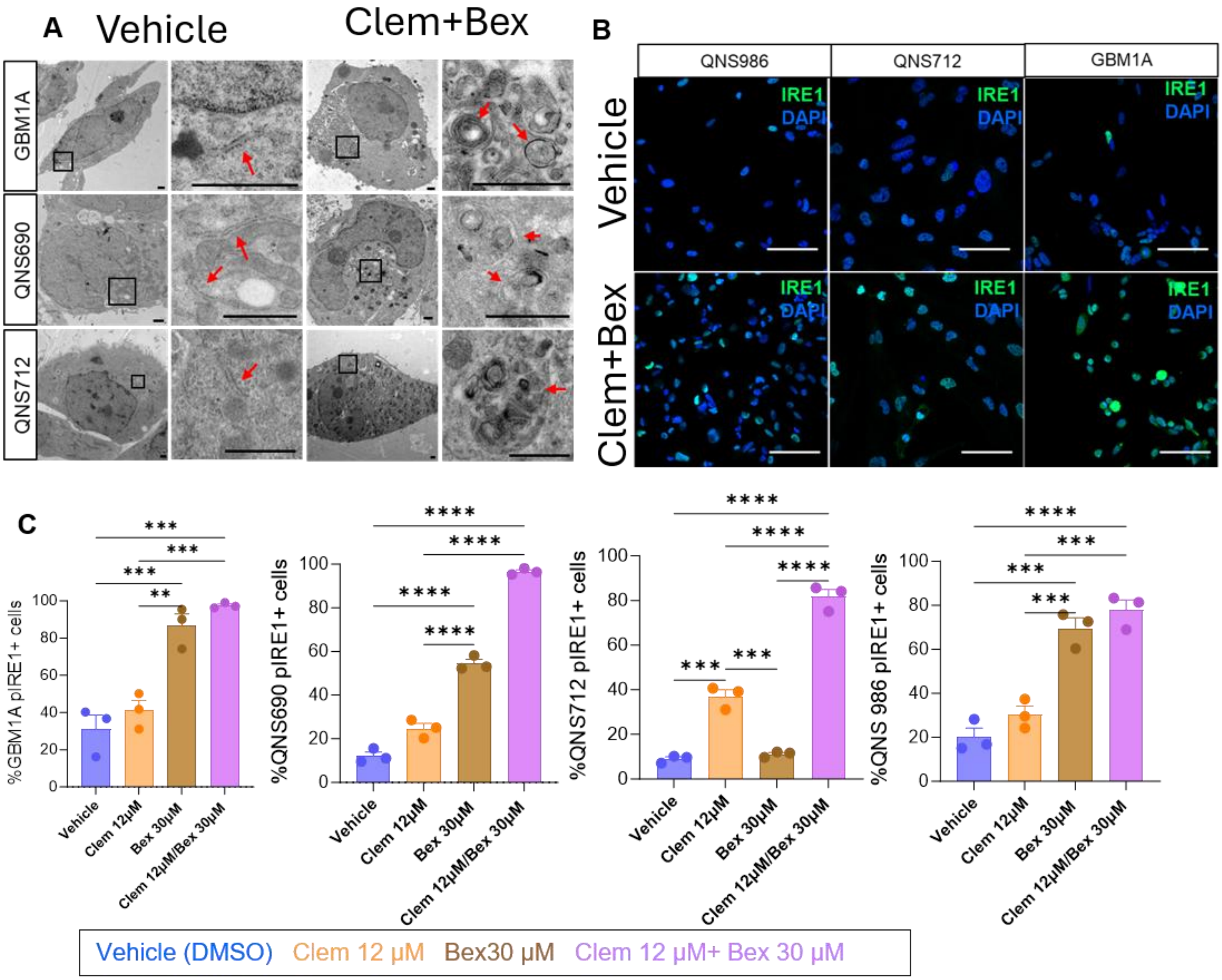


**Supplementary Figure 10. Clemastine and bexarotene induce ER stress and activation of the unfolded protein response in patient-derived GSCs. A**) TEM of GBM1A, QNS690, and QNS712 cells treated with vehicle (DMSO) or Clem+ Bex for 72 h. Combination-treated cells displayed prominent ultrastructural features of ER stress, including dilated endoplasmic reticulum, membrane whorls, and autophagic/autolysosomal structures (red arrows), consistent with activation of ER stress and adaptive stress-response pathways. Scale bars as indicated. **B)** Representative immunofluorescence images of IRE1α (green) in QNS986, QNS712, and GBM1A cells following treatment with vehicle or Clem+Bex. Increased IRE1α expression was observed following treatment, indicating activation of the unfolded protein response (UPR). Scale Bars: 100 μm. **C)** Quantification of IRE1α-positive cells across patient-derived GSC lines treated with vehicle, Clem, Bex, or the combination. Combination treatment produced the strongest induction of IRE1α expression, consistent with robust activation of ER stress signaling. Data are presented as mean ± SEM from independent biological replicates. Statistical significance was determined by one-way ANOVA with multiple-comparison correction. **P < 0.01, ***P < 0.001, **P < 0.0001.

**Supplementary Figure 11**

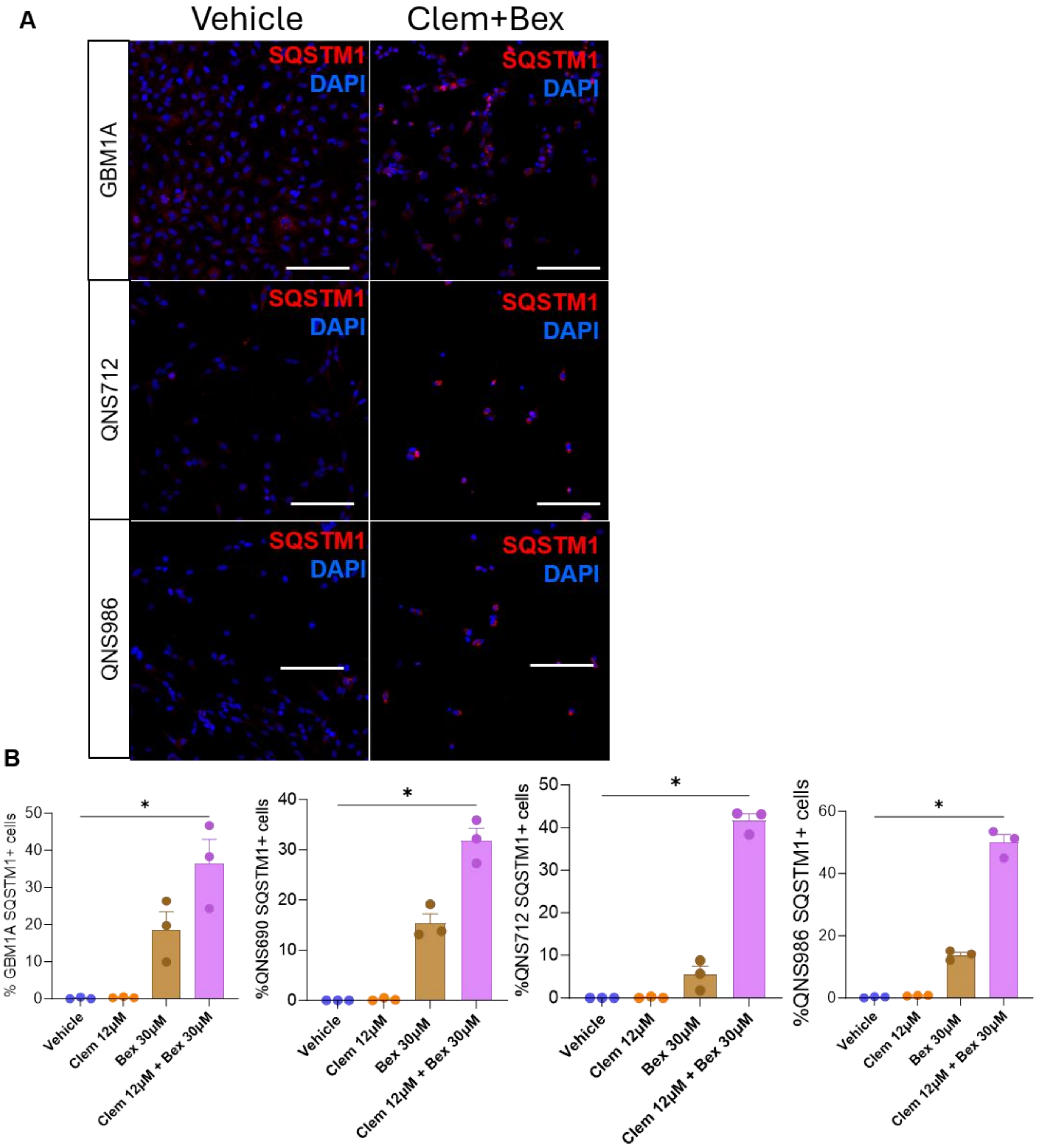


**Supplementary Figure 11. Clemastine–Bexarotene treatment induces SQSTM1/p62 accumulation across multiple patient-derived GBM BTIC lines. A)** Representative immunofluorescence images of SQSTM1/p62 expression in GBM1A, QNS690, QNS712, and QNS986 BTICs treated with vehicle, Clem, Bex, or combination therapy. Scale Bars: 200 μm. **B)** Quantification of SQSTM1/p62-positive cells across patient-derived GBM BTIC lines demonstrating robust induction of autophagy-associated stress responses following treatment. Data are presented as mean ± SEM from independent experiments. Statistical significance was determined using one-way ANOVA with multiple-comparisons correction. *$P < 0.05$.

**Supplementary Figure 12**

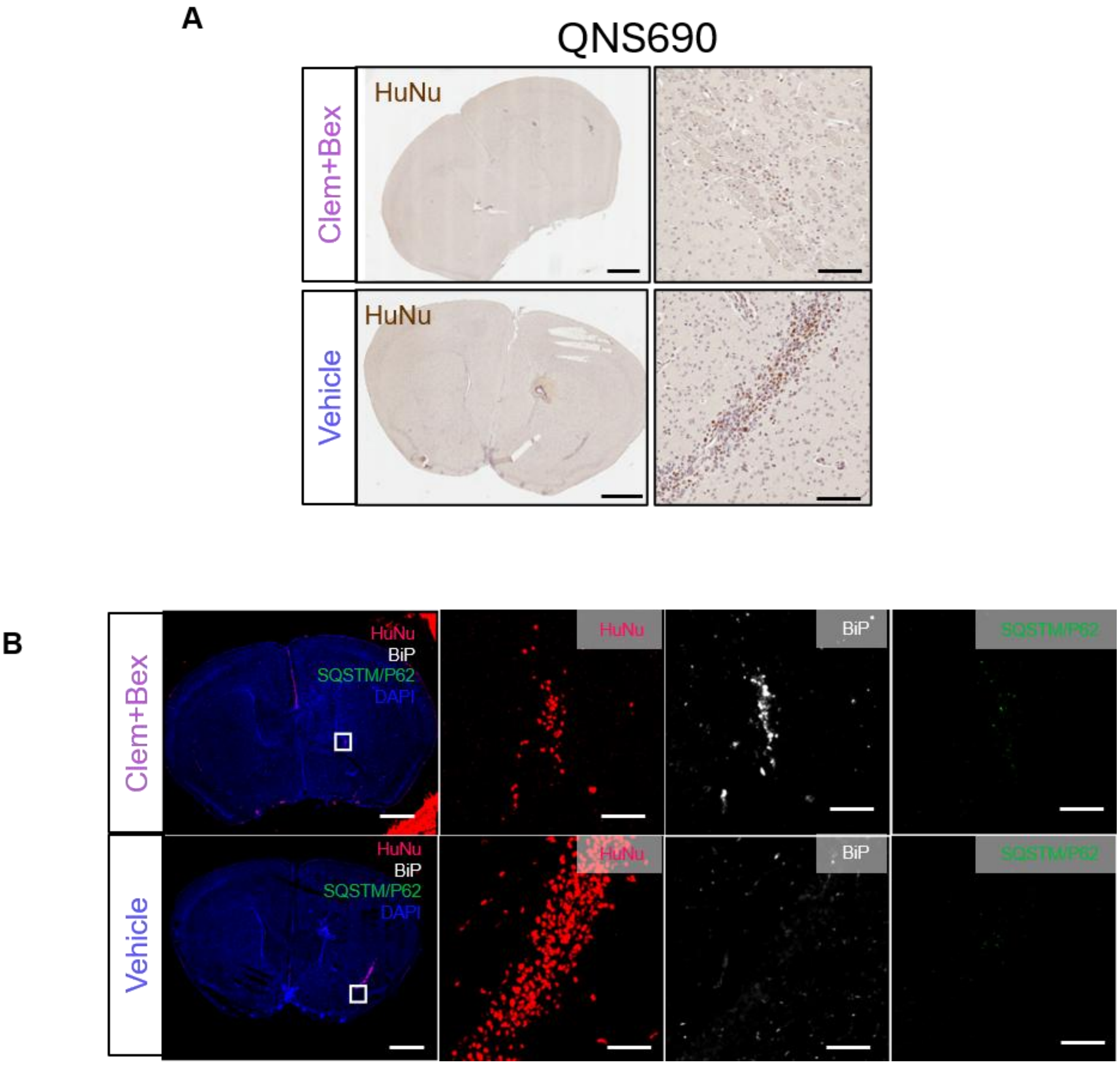


**Supplementary Figure 12. Combined metabolic therapy induces ER stress and autophagy signaling in orthotopic GBM tumors.** **A)** Representative immunofluorescence images of BiP and SQSTM1/p62 expression in orthotopic GBM1A tumors treated with vehicle or Clem+Bex. Scale Bars: 1 mm, 50µm. **B)** RepresentativemIHC images of BiP and SQSTM1/p62 expression in orthotopic QNS690 tumors following treatment. Tumor cells were identified using human nuclei (HuNu) staining. Scale Bars: 1 mm, 50µm.

#### Supplementary Figure 13

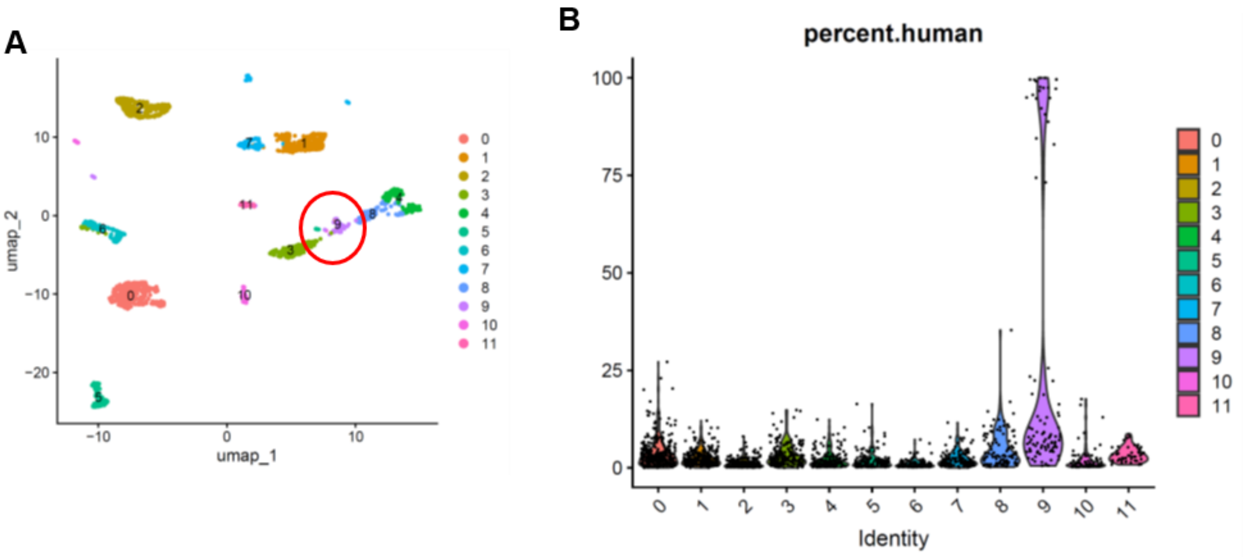


C

| Cluster | Identity | Key marker genes from our data |
|---|---|---|
| 0 | Endothelial | Cldn5, Ptprb, Abcb1a, Slco1a4, Sox18 |
| 1 | Neurons (excitatory-like) | Kcnj6, Kcnh7, Chrm3, Clstn2, Rtn4r |
| 2 | Oligodendrocytes | Sox10, Hapln2, Gpr62, Nipal4 |
| 3 | Microglia | C1qa, C1qb, C1qc, Tyrobp, Ctss, Fcer1g |
| 4 | Neural progenitor / astro-like | Rfx4, Gpc5, Nr2e1, Rgs20 |
| 5 | Pericytes / smooth muscle | Myh11, Aoc3, Cd248, Aspn, Pln |
| 6 | Activated microglia / macrophages | Il1a, C5ar1, Bcl2a1b, Lilrb4a, Nfkbid |
| 7 | OPCs (oligodendrocyte precursors) | Vcan, Myt1, Rgs9, Lhfpl3 |
| 8 | Astrocytes ( doublets) | Aldoc, Aldh1l1, Fgfr3, Cspg5, Cxcl14 |
| 9 | **Human tumor cells (neuronal-like)** | Nrgn, Camk2n1, Cplx2, Map1a, Vamp2 ***(+ high percent.human)*** |
| 10 | Ependymal / ciliated cells | Rsph4a, Dynlrb2, Cfap52, Cfap161 |
| 11 | Oligodendrocytes (myelinating, mature) | Mbp, Mobp, Bcas1, Plekhb1 |

**Supplementary Figure 13. Identification of human GBM cells within the murine tumor microenvironment by single-cell RNA sequencing. A)** UMAP visualization of integrated single-cell RNA-sequencing data from orthotopic glioblastoma tumors. Cluster 9 (red circle) was identified as the putative human tumor population based on elevated human transcript content and expression of neuronal-like glioblastoma markers. **B)** Distribution of human transcript abundance across clusters. Cluster 9 exhibited the highest percentage of human-derived reads, confirming its identity as the human GBMcompartment. **C)** Annotation of major cell populations based on canonical marker gene expression. Identified populations included endothelial cells, neurons, oligodendrocytes, microglia, activated microglia/macrophages, neural progenitor/astro-like cells, pericytes, oligodendrocyte precursor cells (OPCs), astrocytes, ependymal cells, mature oligodendrocytes, and a human neuronal-like tumor cell population. Representative marker genes used for annotation are shown. These analyses enabled separation of human GBM cells from host-derived brain and immune populations for downstream transcriptional analyses.

#### Supplementary Figure 14

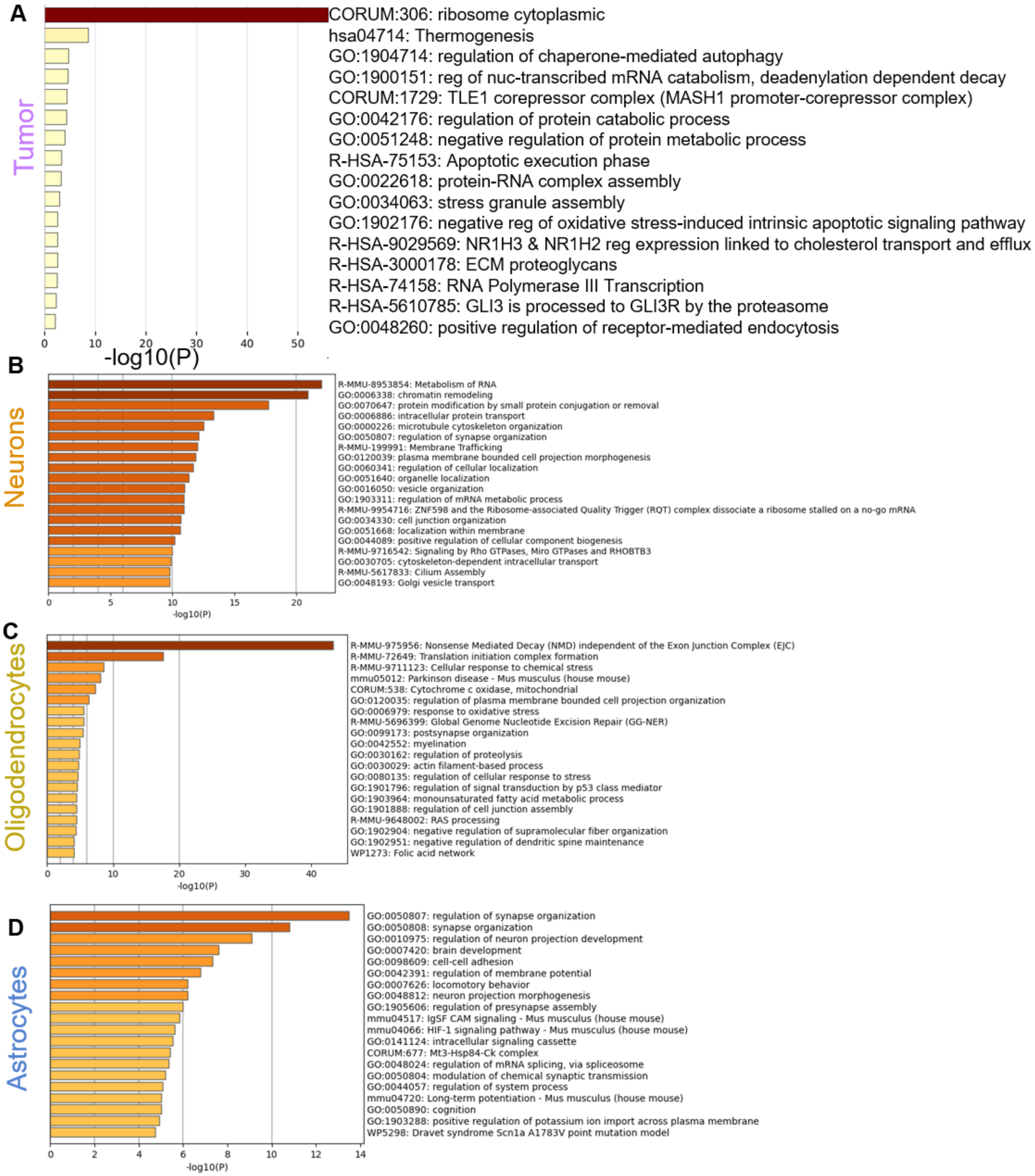


**Supplementary Figure 14. Cell type–specific transcriptional responses to Clemastine+Bexarotene. A–C)** Metascape pathway enrichment analyses of neurons (A), oligodendrocytes (B), and astrocytes (C), showing changes in cellular maintenance, metabolism, and homeostatic pathways. **D)** Pathway enrichment of human GBM cells demonstrating activation of cholesterol dysregulation, ER stress, autophagy, apoptosis, and LXR-associated cholesterol efflux pathways.

**Supplementary Figure 15**

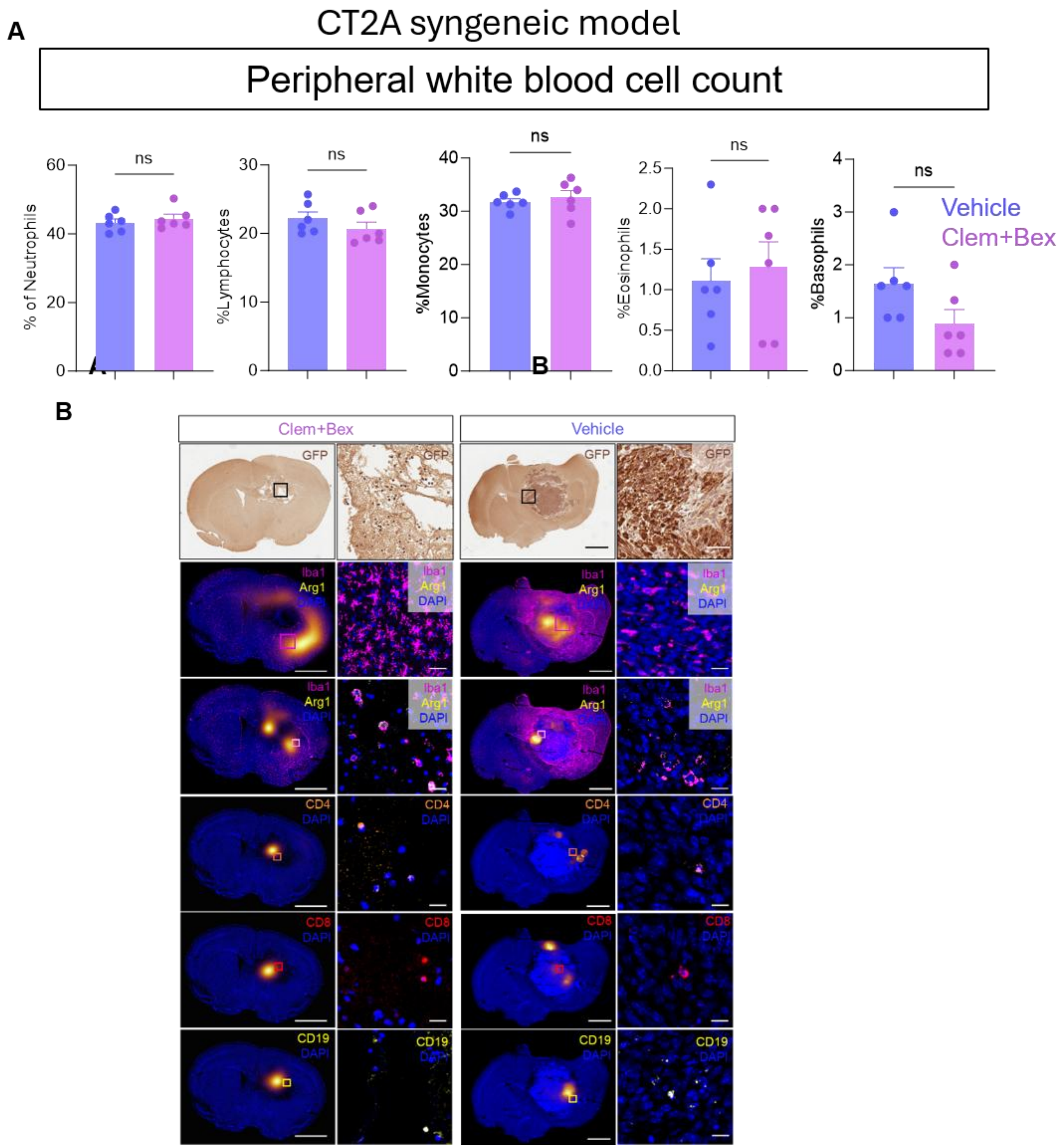


**Supplementary Figure 15. Intracranial clemastine and bexarotene treatment does not induce peripheral immune alterations in a syngeneic GBM model (CT-2A). A)** Peripheral white blood cell analysis of CT2A tumor-bearing mice treated with vehicle or intracranial Clem plus Bex. No significant differences were observed in circulating neutrophils, lymphocytes, monocytes, eosinophils, or basophils, indicating that treatment does not induce major systemic hematologic changes. **B)** Representative histological and IHC analyses of CT2A tumors following treatment. GFP immunohistochemistry revealed altered tumor cell morphology in Clem+Bex-treated mice compared with vehicle controls. mIHC, revealed no significant changes in number of immune cell populations, except for CD4+ T cells consistent with early immune activation. Tissue heatmaps show differential distriburion of macrophages in the tumor. Scale Bars: 1 mm, 25μm

#### Supplementary Figure 16

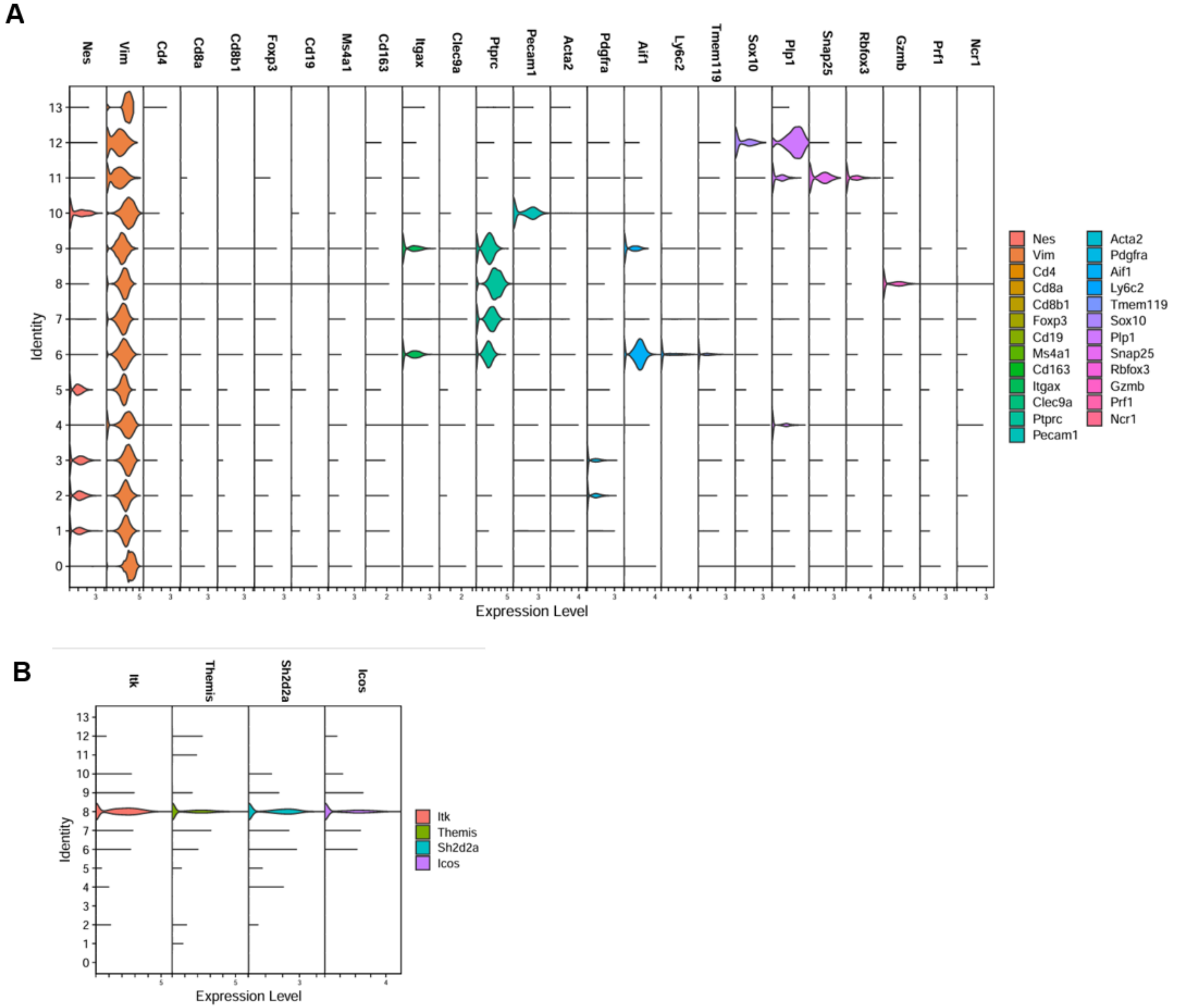


**Supplementary Figure 16. Annotation of cell populations identified by single-cell RNA sequencing in CT-2A tumors. A)** Violin plots showing expression of canonical marker genes used for cluster annotation. Tumor populations were identified by expression of *Nes* and *Vim*, whereas immune and brain-resident populations were annotated using established lineage markers. **B)** Expression of lymphocyte-associated markers supporting identification of T-cell and activated immune populations. Clusters were subsequently classified as tumor cells, endothelial cells, activated macrophages, inflammatory macrophages, astrocytes, pericytes, oligodendrocytes, NK cells, and T lymphocytes.

**Supplementary Figure 17**

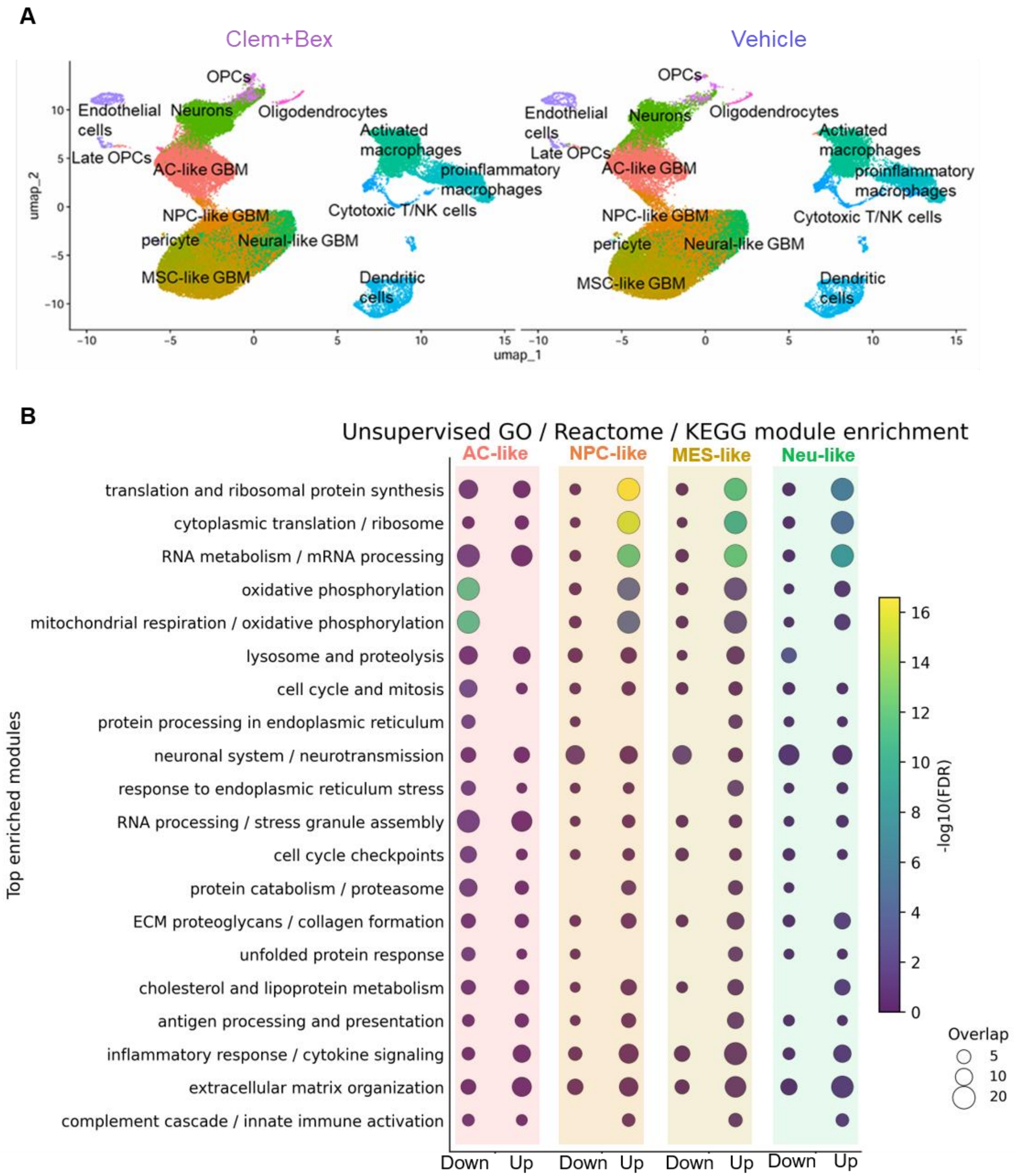


**Supplementary Figure 17. Conserved transcriptional responses to Clem+Bex across GBM cell states. A)** UMAP of single-cell RNA-seq from vehicle- and Clem+Bex-treated CT-2A tumors showing major brain, immune, and tumor cell populations, including AC-, NPC-, MES-, and Neu-like GBM states. **B)** GO, Reactome, and KEGG pathway enrichment analyses demonstrating conserved activation of cholesterol dysregulation, ER stress, UPR, proteotoxic stress, inflammatory signaling, and innate immune pathways across all tumor states.

#### Supplementary Figure 18

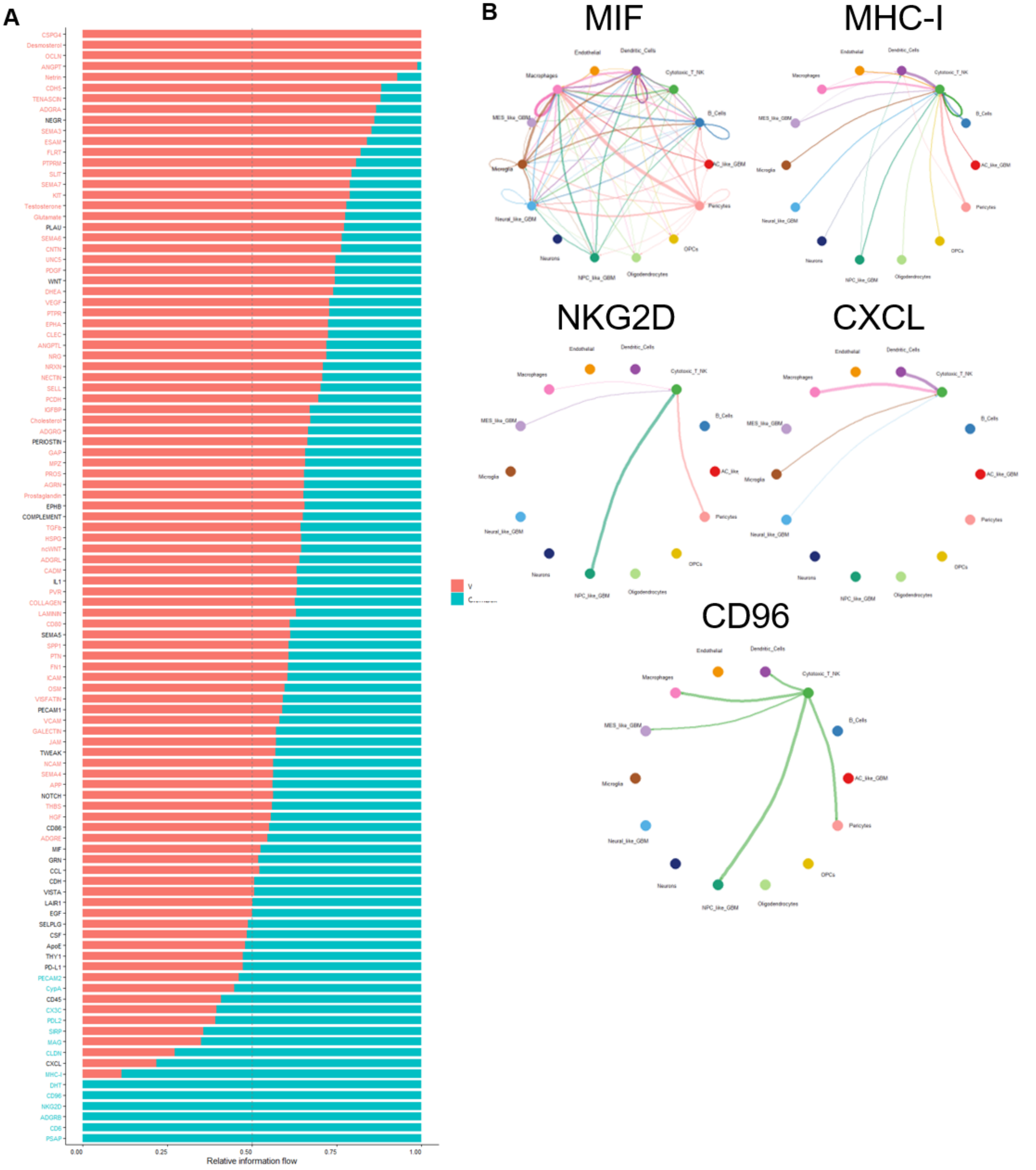


**Supplementary Figure 18. Clemastine+Bexarotene remodels cell–cell communication toward immune surveillance. A)** CellChat rankNet analysis comparing signaling pathway activity between vehicle- and Clem+Bex-treated tumors. **B)** CellChat networks for MIF, MHC-I, NKG2D, CXCL, and CD96 signaling, demonstrating enhanced antigen presentation, chemokine signaling, and cytotoxic immune communication following treatment.

#### Table S1

| Clinical, Radiographic, and Pathologic Characteristics of Patient-Derived Glioblastoma Models | | | | | | | | | | | | | |
|---|---|---|---|---|---|---|---|---|---|---|---|---|---|
| Clinical Information | | | | Pre-Operative MRI Results | | Pathology Results | | Molecular Characteristics | | | | | |
| Subject ID | Age (yr)/ Sex | Primary/ Recurrent | Prior Therapy (before surgery) | T1 Post-Contrast (axial) | T2/FLAIR (axial) | H&E (40x) | Integrated Dx | IDH Status | MGMT Promoter Methylation | TP53 Mutation | EGFR Amplification | ATRX Expression | TERT Promoter Mutation |
| QNS690 | 70/M | Primary | None | 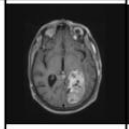 | 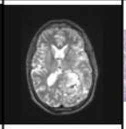 | 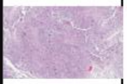 | Left brain tumor: Glioblastoma, IDH-wildtype (WHO grade IV) | Wildtype | Unmethylated when newly dx (at recurrence, path came back at methylated) | Yes | No | Retained | Yes |
| QNS712 | 74/M | Primary | None | 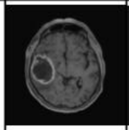 | 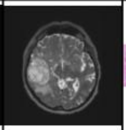 | 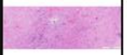 | Right brain tumor: Glioblastoma, IDH-wildtype (WHO grade IV) | Wildtype | Unmethylated | Yes | No | Retained | Yes |
| QNS986 | 50/F | Primary | None | 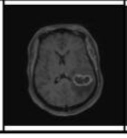 | 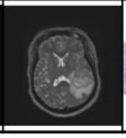 | 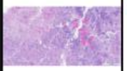 | Left brain tumor: Glioblastoma, IDH-wildtype (WHO grade IV) | Wildtype | Methylated | Yes | Yes | Retained | Yes |

**Supplementary table 1. Clinical, radiographic, histopathological, and molecular characteristics of patient-derived GSC models used in this study.** Clinical information, preoperative imaging findings, histopathological features, and molecular characteristics of the patient-derived GSC models QNS690, QNS712, and QNS986 established from primary glioblastoma specimens obtained at Mayo Clinic through the BRIDGE biobank. Histological diagnoses were assigned according to the WHO Classification of Tumors of the Central Nervous System. Molecular features include IDH status, MGMT promoter methylation, TP53 mutation status, EGFR amplification, ATRX expression, and TERT promoter mutation status. Representative preoperative magnetic resonance imaging (MRI) and H&E staining are shown when available. The GBM1A (Patient 1) model used throughout this study has been previously described and characterized by Vescovi and colleagues56 and is therefore not included in this table.
